# A multimodal approach for visualization and identification of electrophysiological cell types *in vivo*

**DOI:** 10.1101/2025.07.24.666654

**Authors:** Eric Kenji Lee, Asım Emre Gül, Greggory Heller, Anna Lakunina, Han Yu, Andrew Shelton, Shawn Olsen, Nicholas A. Steinmetz, Cole Hurwitz, Santiago Jaramillo, Pawel F. Przytycki, Chandramouli Chandrasekaran

## Abstract

Neurons of different types perform diverse computations and coordinate their activity during sensation, perception, and action. While electrophysiological recordings can measure the activity of many neurons simultaneously, identifying cell types during these experiments remains difficult. To identify cell types, we developed PhysMAP, a framework that weighs multiple electrophysiological modalities simultaneously to obtain interpretable multimodal representations. We apply PhysMAP to seven datasets and demonstrate that these multimodal representations are better aligned with known transcriptomically-defined cell types than any single modality alone. We then show that such alignment allows PhysMAP to better identify putative cell types in the absence of ground truth. We also demonstrate how annotated datasets can be used to infer multiple cell types simultaneously in unannotated datasets and show that the properties of inferred types are consistent with the known properties of these cell types. Finally, we provide a first-of-its-kind demonstration of how PhysMAP can help understand how multiple cell types interact to drive circuit dynamics. Collectively, these results demonstrate that multimodal representations from PhysMAP enable the study of multiple cell types simultaneously, thus providing insight into neural circuit dynamics.

## Introduction

Single cell transcriptomics, electrophysiology, and morphological reconstruction have identified numerous neuronal cell types in the brain^1–3^. These cell types, with their specific patterns of gene expression, morphology, and connectivity^4^, form microcircuits that perform neural computations and drive behavior^5–7^. For instance, in mice, interactions between excitatory pyramidal, parvalbumin-positive (PV^+^), somatostatin-positive (SOM^+^), and vasoactive intestinal peptide-positive (VIP^+^) neurons enable computations such as gain modulation^8^ and selective enhancement of salient visual stimuli^9^. Despite such demonstrations, only one cell type can be monitored simultaneously in most experiments, thus precluding a full understanding of how cell types contribute to the brain’s many computations^10^. Moreover, these techniques are infeasible or technically challenging in species other than mice and regions other than neocortex because of a lack of effective viral constructs^11,12^; these are in addition to barriers to optical access^13–15^. These barriers prevent the study of cell types in deep structures and in organisms most relevant for understanding human psychiatric disorders, such as primates.

In contrast, high-density extracellular electrophysiological recordings can simultaneously sample from numerous neurons throughout the brain without genetic or optical access^16–20^, including in humans^21,22^. Yet the cellular identities of recorded neurons remain largely invisible since the correspondence between cell types and their *in vivo* electrophysiological properties remains poorly understood, thus preventing a comprehensive understanding of circuit dynamics. One way to mitigate this drawback is to develop an analysis approach that can delineate cell types in electrophysiological recordings *in vivo*. Ideally, such an approach should have the following capabilities: First, this approach should be able to maximally incorporate electrophysiological information to identify cell types by combining multiple modalities together. Numerous studies suggest that waveform shape^23–26^, inter-spike interval (ISI) distributions^27–29^, autocorrelogram (ACG)^30^, and peri-stimulus time histograms (PSTHs)^31–33^ are potentially informative about cell type. However, which modalities to combine (and how to best combine them) is not well understood. Second, cell types are not discrete as they often lie along a continuum of function^34^. Thus, any approach to delineate cell types from electrophysiological data should be able to visualize and explore the heterogeneity of cell types in an interpretable manner (e.g., how biological features vary within a cell type). Third, in the cases where cell type identity is available, representations should logically cluster according to ground truth. Such clustering would provide confidence that this structure can be used to identify cell types in other datasets not containing ground truth. Fourth, the approach should be usable in real use cases to identify cell types in new data using existing ground truth after ensuring that the two datasets are well matched. Finally, this method should be able to excel at all these things while remaining performant enough to iterate quickly over different modalities and datasets.

Here we present “PhysMAP,” an approach satisfying all of these capabilities. PhysMAP leverages advances in multimodal representation learning from computational biology^35,36^, and combines non-linear dimensionality reduction^37^ with a weighted-nearest neighbor graph (WNN)^38^. Through this approach, PhysMAP individually weighs multiple electrophysiological modalities on a per-unit basis to create an interpretable multimodal representation for unsupervised cell type identification from electrophysiological recordings. We first apply PhysMAP to seven datasets from different neuroscience labs and show that it: 1) optimally combines modalities and identifies cell types when ground truth is available, 2) can be used as a tool to explore and cluster putative cell types without annotations, 3) identifies cell types in datasets with partial high-quality annotations, 4) detects batch effects when present, and 5) runs on a laptop and provides results in a few minutes. To showcase the real-world usefulness of our approach, we identify cell types in a dataset with incomplete ground truth and then assess the properties of several cell types interacting simultaneously during behavior. PhysMAP thus provides a multi-faceted approach for identifying cell types in electrophysiological recordings of behaving animals, advancing our ability to study simultaneous circuit dynamics of multiple neuronal types during behavior.

## Results

### PhysMAP combines multiple electrophysiological modalities to uncover cell type and laminar structure

*Ex vivo* experiments have shown that the relationship between cell type and electrophysiological signature spans multiple properties^2,39,40^, suggesting that cell types could be identified from *in vivo* extracellular recordings. We examined our first desired property of a multimodal tool: whether combining multiple electrophysiological modalities could better align with cell types than any single modality. PhysMAP unifies different modalities by linearly combining high-dimensional UMAP graphs using a weighted nearest neighbor approach^38^ on an arbitrary number of modalities^41^ (see Appendix A for details).

We applied PhysMAP to *in vivo* juxtacellular recordings from the mouse somatosensory cortex, which provided low-noise somatic waveforms and ground truth cell type information (n = 246) from eight cell types^32^. UMAP applied to waveform shape^25,42^ produced 2D embeddings that aligned well with underlying cell types and laminar structure (Fig. 1A), though some cell types (like layer 4 excitatory cells [E-4]) appeared split. In contrast, this split disappeared in the ISI modality (Fig. 1B), but there was less clear delineation between PV^+^ and SOM^+^ cells. However, E-4 cells were well separated from E-5 (layer 5 excitatory cells) cells and other cell types. These complementary views suggest that optimally combining multiple modalities could provide more robust cell type delineation, as unsupervised multimodal approaches benefit when signals correlate across modalities while noise remains uncorrelated^43^.

**Figure 1.**
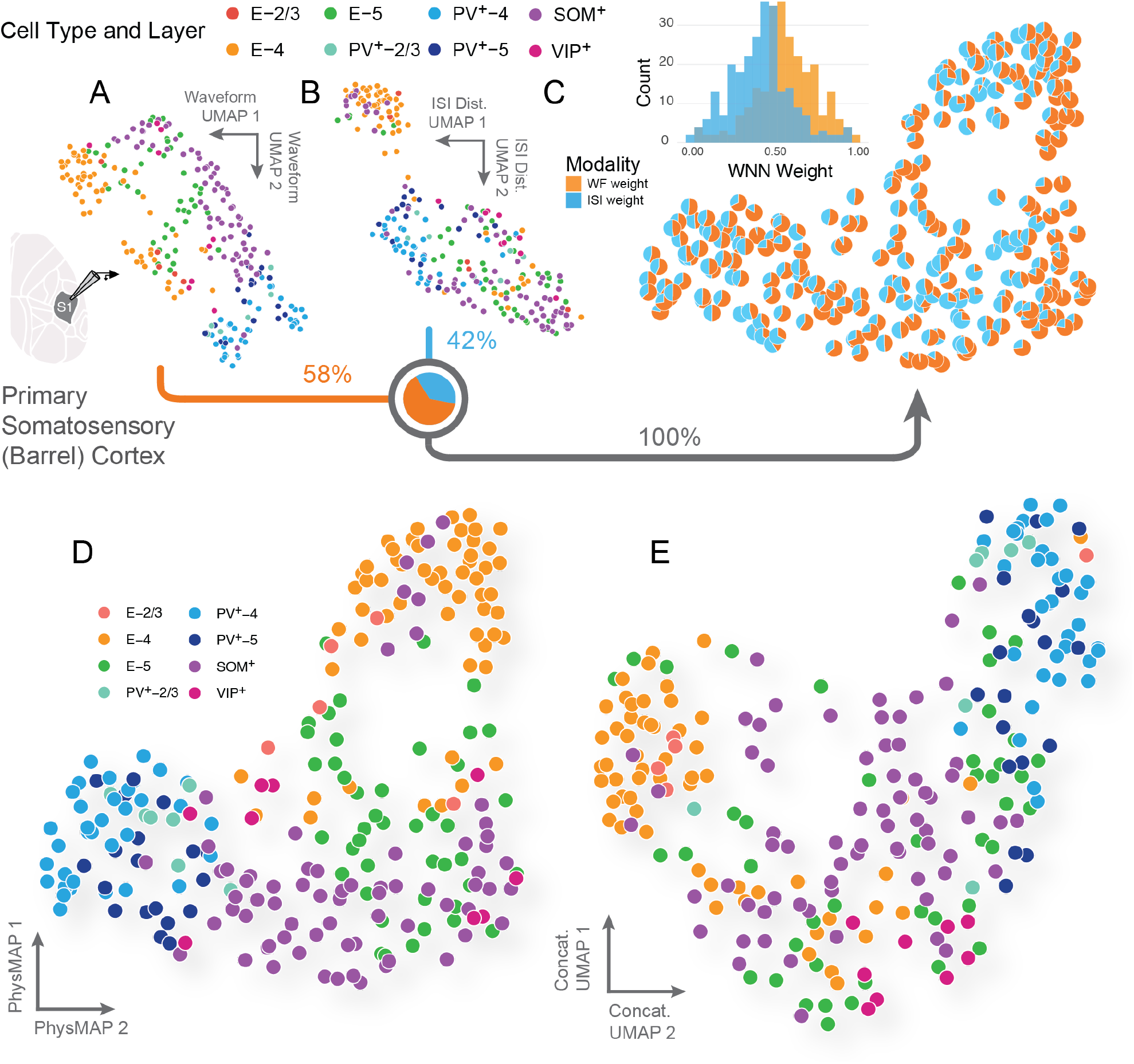
Multi-modal combination of physiological properties leads to structured representations that align with cell types. (**A**) UMAP on normalized average waveform shapes of neurons (n = 246) recorded *in vivo* juxtacellularly from mouse primary somatosensory cortex^32^. Each unit is colored according to one of eight ground truth cell types. Each modality was combined in different proportion according to the sum of each modality’s per-unit contribution to the total; this total contribution is listed as a percentage below each scatter plot. (**B**) UMAP on ISI distributions of the same neurons in (A). (**C**) Each neuron in PhysMAP’s WNN representation weighs each modality differently. These modal contributions are the sum of nearest neighbor edge modality weights (*β*_Modality_ in Fig. A.13J) and are shown in their proportion of the per-unit WNN weight (pie charts) or across all units (inset histogram). (**D**) PhysMAP’s combined multimodal representation obtained using the WNN approach^38^. Again, each neuron is colored according to their ground truth cell type and layer. (**E**) Both modalities were concatenated into a single data vector per unit and passed into UMAP. This represents each modality in unweighted combination (as opposed to the WNN approach).

We examined how PhysMAP weights each modality per-neuron (Fig. 1C). For most units, waveform shape was most informative (58% vs. 42% for ISI)^29^, though certain regions of the projection showed greater ISI importance. The multimodal representation obtained by weighting each modality per-neuron appeared to best separate cell types visually (Fig. 1D). This visualization revealed that PV^+^ cells cluster tightly and segregate by layer, while SOM^+^ cells occupy a larger area, perhaps a result of underlying physiological diversity^44–46^.

To test whether PhysMAP’s improvements were a trivial consequence of including more information, we applied UMAP to a single feature vector (unweighted combination of all modalities) by concatenating both waveform shape and ISI (Fig. 1E). The resulting representation showed a weaker structure than PhysMAP (compare Fig. 1E to Fig. 1D), with poorer clustering of E-5, SOM^+^, and E-4 neurons. These results suggest PhysMAP’s weighted averaging approach better utilizes multiple modalities.

We also replicated this analysis with three modalities by adding peri-stimulus time histograms (PSTHs) aligned to whisker deflection (Fig. S1A, B, and C, respectively). PSTH could also separate cell types, but was not the most informative modality for any units (Fig. S1D). Again, the three-modality PhysMAP representation showed better clustering of PV^+^ and E-4 cells visually (Fig. S1E vs. in Fig. 1D) while the concatenated three-modality representation remained poorly clustered (Fig. S1F) again reinforcing that naïve combination doesn’t improve representation quality. Since PSTH contributed minimally and its inclusion would preclude assessment of other functional properties due to analytic circularity^47^, we excluded it from subsequent analyses.

*These results show that PhysMAP satisfies the first requirement of an ideal tool: it can combine multiple modalities to obtain a representation that is superior to any single modality*.

### Interpretable representations from PhysMAP enable biological insight

Even when cell types have been identified through optotagging, they often encompass multiple subclasses^44^. In other words, cell types are rarely discrete, with physiological variation within a class being the norm^34^. Thus, a second desired property of any tool to delineate cell types from electrophysiological data is that it should also be able to visualize physiological diversity within and across defined types.

Here we demonstrate that PhysMAP’s visualizations excel at this purpose. Fig. 2A shows the PhysMAP plot from Fig. 1D with marker radius scaled to reflect waveform spike width, peak-to-trough ratio, and onset latency. Examining spike width (Fig. 2A, left), clear patterns emerge: PV^+^ cells are uniformly narrow regardless of layer; SOM^+^ cells span a gradient of widths overlapping with E-5 cells at their widest; and E-4 cells have the widest spikes. This heterogeneity in SOM^+^ cells likely reflects several subtypes with varying widths^50^ and distinct computational roles^45^. Waveform peak-to-trough ratios also differ between cell types (Fig. 2A, center) with PV^+^ and SOM^+^ cells having the smallest ratios and both E-4 and E-5 cells occupying the largest ratios. Firing rate onset latency (Fig. 2A, right) is fastest for PV^+^ cells and slowest for SOM^+^ cells, consistent with^32^. These interpretations also held when the trial-averaged firing rate response to a whisker deflection (peri-stimulus time histogram, PSTH) was included (Fig. S4A). We note that PSTH was excluded here to avoid analytic circularity^47^ as, in the next section, we examine the PSTH of each unsupervised cluster.

**Figure 2.**
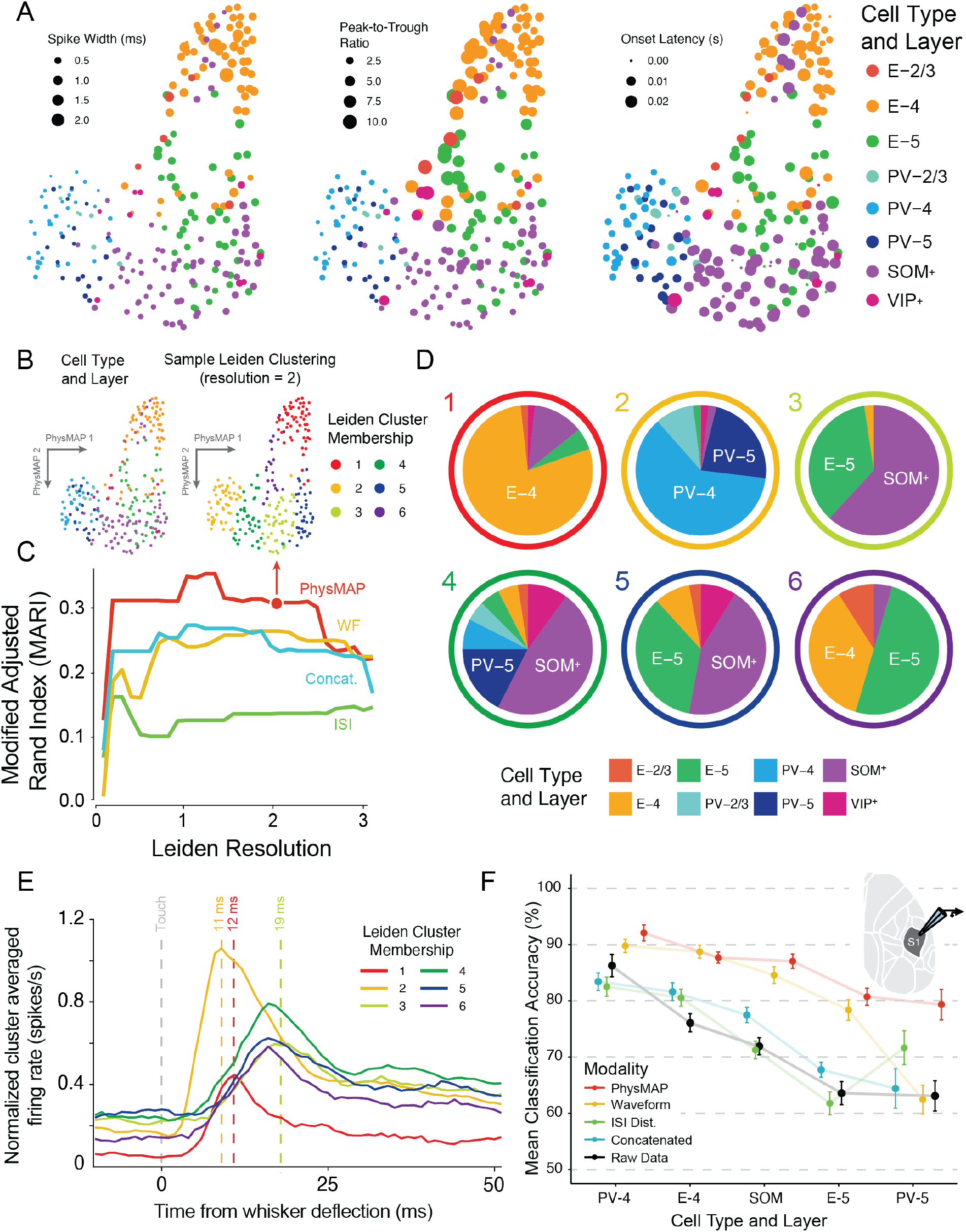
PhysMAP identifies cell types from juxtacellular recordings better than any modality alone. **(A)** Juxtacellular single unit recordings (n = 246) from mouse primary somatosensory cortex were collected under two modalities: juxtacellular waveform shape and inter-spike interval distribution in response to whisker deflection. Shown are two-dimensional PhysMAP visualizations of these units. Each unit is colored according to its ground truth cell type as deduced by optogenetic tagging in addition to layer information. The spike width (time from trough to peak), peak-to-trough ratio (ratio of trough and peak absolute amplitude), and spiking onset latency (trial-averaged time to half-peak in firing rate) for each neuron. Marker sizes shown are proportional in area to the log of the scaler value. (**B**) The PhysMAP projection of neurons with their ground truth cell type and layer (**left**) next to an example Leiden clustering^48^ with resolution parameter set to 2 (**right**). (**C**) Leiden clustering is applied to the UMAP graphs of each modality alone (waveform [WF] or ISI dist. [ISI]), unweighted combination (concat.), or in weighted (PhysMAP) and shown with the associated modified adjusted Rand index (MARI;^49^) across a range of resolution parameter values. Arrow marker indicates the clustering on PhysMAP used above in (B). Waveform metrics and “raw data” (that is, without constructing a UMAP graph and projecting it) were omitted in this analysis because these do not yield a high-dimensional UMAP graph. (**D**) Each of the Leiden clusters located in (B) are shown broken down into their constituent cell types. Outer ring color indicates the Leiden cluster and inner pie chart denotes the relative proportions of underlying cell types within said Leiden cluster. (**E**) Each of the Leiden clusters in (B, right) and (D) are shown with their respective normalized and smoothed peri-stimulus time histograms aligned to a whisker deflection (“touch” stimulus). Also labeled are the firing rate peak times of several Leiden clusters relative to the stimulus (dashed lines). (**F**) A gradient boosted tree model (GBM) classifier was trained on this multimodal data with 5-fold cross-validation on each modality’s UMAP graph individually or on the multimodal WNN graph. The same classifier was also trained directly on the “raw data” (that is, without constructing a UMAP graph and projecting it; black line). The balanced accuracy performance of this classifier (mean *±* S.E.M.) on held-out data for each modality or the combined modalities for the five cell type classes with over 10 samples (units) each.

These insights are not apparent when examining metrics without the PhysMAP representation. A scatter plot of spike width versus peak-to-trough ratio (Fig. S2) shows some structure but appears as one continuous relationship, failing to clearly separate SOM^+^, PV^+^ and excitatory cells. Histograms of these metrics show considerable overlap between cell types (Fig. S2B).

*Thus, PhysMAP satisfies the second property of an ideal tool as it provides an interpretable visualization that effectively combines multimodal electrophysiological data, facilitating exploration of physiological diversity while maintaining the relationship to underlying cell types*.

### Combining multiple modalities improves unsupervised clustering

There are many cases where annotations are not available, e.g. in electrophysiological datasets without optotagging. In such cases it is often desirable to uncover cell types, via unsupervised methods^1,2,21,23,24,26,29,51^. Thus, the third property of a tool to delineate cell types from electrophysiological data is that it should logically group and cluster cell types within unlabeled datasets. To do this, we evaluated whether PhysMAP representations align with underlying molecular cell types in a fully unsupervised analysis, asking the question: “without access to ground truth, how well do unsupervised methods identify cell types?”

We assessed cluster-to-ground-truth alignment using the modified adjusted Rand index (MARI;^49^), to estimate if ground truth cell types align with individual unsupervised clusters. We applied Leiden clustering to the high-dimensional graphs produced by PhysMAP and each individual modality and then calculated MARI scores between clustering results and ground truth labels across a range of resolution parameters (cluster size priors; see Methods: MARI Calculation for details). Fig. 2B shows neurons in PhysMAP colored by cell type (top left) alongside a sample Leiden clustering with resolution set to 2 (top right) which produced the same number of clusters as prominent cell types. PhysMAP produced higher MARI scores than any individual modality or unweighted combination across all Leiden resolutions (Fig. 2C), indicating superior unsupervised capture of cell type structures at all levels of granularity (see Fig. S3 for clustering at other resolutions). This result also held when PSTH was included as a modality (Fig. S4B). Choosing a resolution of 2 (the maximum value before MARI dropped), six clusters were produced with varying proportions of cell types within each (Fig. 2D). We found that while not all clusters captured single cell types, many showed high specificity: E-4 cells dominated cluster 1, PV^+^ cells (especially PV-4) dominated cluster 2, and SOM^+^ cells were substantially represented in cluster 3. When a traditional clustering approach (a Gaussian mixture model) was used on spike waveform metrics Fig. S5B, this resulted in greater mixing of cell types within each unsupervised GMM cluster (Fig. S5C).

To assess the functional properties of these unsupervised clusters, we examined their responses to whisker deflection stimuli (Fig. 2E). Several notable features aligned with previous findings in the literature: cluster 2 (predominantly PV^+^) showed large, low-latency (11 ms) firing rate increases, matching properties of PV^+^ cells reported by^32^; cluster 1 (predominantly E-4) responded with similar latency but much smaller firing rate changes consistent with excitatory cells receiving direct thalamo-cortical input; and cluster 3 (containing many SOM^+^ cells) was the slowest responding, consistent with previous characterizations of SOM^+^ cells. While SOM^+^ cells were split among several clusters, this is unsurprising given their diversity^44,45^. Traditional waveform metrics similarly struggled to cluster SOM^+^ cells, as shown by their distribution split across separate GMM clusters (Fig. S2B and C).

*These results show that PhysMAP has another property desired of an ideal tool—provide unsupervised clustering methods an excellent opportunity to capture underlying cell types, even when annotations are unavailable*.

### PhysMAP delineates cell types better than any modality alone or in unweighted combination

The previous section showed that PhysMAP’s multimodal representations align better with underlying cell types from an unsupervised perspective. However, there are also datasets that contain cell type information; do representations continue to identify these cell types assessed via a classifier trained on PhysMAP’s structure? We now examine whether these advantages persist in a supervised analysis by training a classification algorithm on PhysMAP’s representation and seeing if it outperformed other classifiers trained on individual modalities or in unweighted combinations. In this way, we examine PhysMAP’s performance on datasets that both completely lack ground truth or are also fully characterized.

We trained classifiers on high-dimensional graphs to identify cell types using individual modality representations, unweighted concatenation, and PhysMAP. For comparison, we also trained classifiers on derived electrophysiological metrics and raw data without dimensionality reduction. Using a 15-85% test-train split, we trained gradient-boosted tree models (GBM) with five-fold cross-validation on high-dimensional embeddings to identify the five cell types with over 10 examples each (so that at least one data point from each class could be included in the test set; see Methods: Classifier Analysis for details).

PhysMAP matched or exceeded the performance of all other modalities, whether individual or in unweighted combination, across all cell types (Fig. 2F). Waveform shape performed nearly as well as PhysMAP for PV-4, E-4, and SOM^+^ cells, confirming our hypothesis that waveform shape is the most cell type-informative modality (Fig. 1C) and validating our previous findings with WaveMAP^24,25^. The benefit of additional modalities was most evident in improved classification of E-5 and PV-5 neurons.

PhysMAP also demonstrated better classification performance than the modalities in an unweighted combination (concatenated) for all cell types. PhysMAP and waveform shape far outperformed traditional derived waveform metrics (spike width and peak-to-trough ratio; Fig. S4C, gray dashed line), confirming that hand-derived metrics lose important cell type-distinguishing information. Notably, PhysMAP performed better than classifiers trained on raw data, despite the dimensionality reduction step: the GBM classifier trained directly on full-dimensional data from all modalities underperformed both PhysMAP and waveform shape alone (Fig. 1C, black dashed line). While seemingly counterintuitive, this reflects how dimensionality reduction helps “denoise” latent clusters in smaller datasets. Moreover, training directly on raw data sacrifices interpretability, providing only class labels and probabilities without the visualization benefits of PhysMAP.

These results were robust to choices of input dimensionality and classifier type. Classification accuracy patterns remained consistent across embedding dimensions from high (30-dim.) down to low (2-dim.; Fig. S5A) and across classifier types (Fig. S5B). This preservation of performance trends is likely because the input modalities are themselves not very high-dimensional. Principal component analysis showed that the intrinsic dimensionality of each dataset was relatively low, with “elbows” in the scree plots at the third, fourth, and third PC’s representing 91%, 80%, and 93% of total variance for waveform shape, ISI distribution, and PSTH respectively (Fig. S6).

*Thus, PhysMAP satisfies our third property of an ideal cell type identification tool as it intelligently combines multiple modalities to draw out cell type structure better than any modality alone or in unweighted combination, while providing valuable visualizations for exploring physiological diversity and maintaining high classification performance*.

### PhysMAP identifies cell types in extracellular electrophysiological recordings

The juxtacellular recordings analyzed previously represent a best-case scenario for extracellular electrophysiology: these datasets provide unattenuated somatic waveforms, unambiguous single-unit isolation, and preferential targeting of specific cell types. We next asked whether PhysMAP could identify cell types in extracellular probe recordings, which introduce challenges including attenuation of signals by neuropil, poorer single-unit isolation, and the introduction of non-somatic waveforms—this all contributes to increased ambiguity of cell type properties. We analyzed an extracellular dataset from mouse primary auditory cortex with optotagged cells.

In this study, researchers recorded from neurons in A1 (n = 373) with extracellular probes while mice were presented with auditory stimuli. They identified both SOM^+^ and PV^+^ units via optogenetic tagging and recorded each unit’s waveform shape and ISI distribution. When PhysMAP was applied to this data, visualization shows clear separation of both SOM^+^ and PV^+^ cell types (purple and teal, respectively) from untagged, presumably excitatory cells (gray; Fig. 3A).

**Figure 3.**
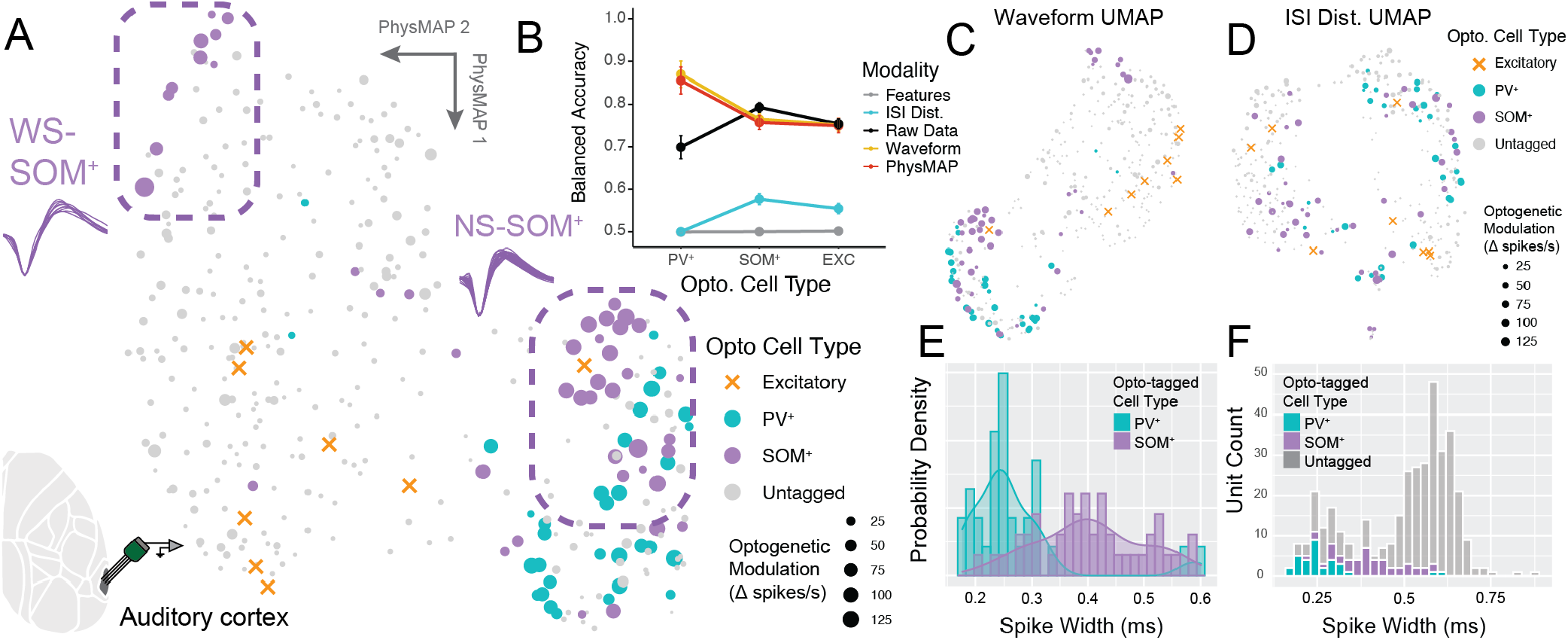
PhysMAP identifies cell types in extracellular recordings. (**A**) PhysMAP applied to putative single unit (n = 373) extracellular recordings from mouse auditory cortex^52^ using silicon multi-channel probes. We used waveform shape and ISI distribution as our modalities and neurons are colored according to cell type whose identities were obtained via optogenetic tagging. Marker size is set according to the increase in spike rate from optogenetic stimulation relative to baseline. Highlighted are a population of wide-spiking SOM^+^ cells (**top**) and narrow-spiking SOM^+^ cells (**right**). Excitatory cells are shown as orange x’s because optogenetic modulation was negative and too small to see by size. (**B**) A GBM classifier with five-fold cross-validation was trained on the 10-dimensional graph of each modality or PhysMAP. The classifier was also trained directly on the raw (un-dimensionality reduced) multimodal data or simply waveform spike width to identify each optotagged cell type or untagged label. The balanced classification accuracy is shown (mean *±* S.E.M.) with many error bars smaller than marker size. (**C**) Normalized average waveforms in this dataset are passed into UMAP and their projection shown with optotagged cells colored and degree of optogenetic modulation (average change in spikes/s during stimulation epochs) shown by marker size. (**D**) Similarly, ISI distributions for these same neurons are also passed into UMAP and their projection shown. (**E**) Gaussian kernel probability density estimates for the distribution of each optotagged type across a range of spike widths is shown both as a histogram and with a kernel density estimator (solid line and shaded region). (**F**) Additionally, spike width is shown for all cells, including untagged ones.

We quantified this separation by training a classifier to identify each cell type using PhysMAP and compared it against classifiers trained on individual modalities or waveform metrics. PhysMAP outperformed ISI distributions, raw data, and waveform metrics but matched performance on waveform shape for PV^+^, SOM^+^, and excitatory cells (Fig. 3B). This equivalence likely occurs because waveform shape is highly informative of underlying cell types, with median per-unit weights of 0.71 for waveform shape versus 0.29 for ISI distribution. This is a demonstration of PhysMAP’s ability to focus only on informative modalities and this prioritization of waveform shape is evident in the visualizations: PhysMAP’s projection closely resembles the waveform shape visualization (Fig. 3C), while the same for ISI distribution shows far less structure (Fig. 3D).

The PhysMAP approach also revealed interpretable cell type heterogeneity. We identified two sub-populations of SOM^+^ cells not documented in the original study: wide-spiking SOM^+^ (WS-SOM^+^, Fig. 3A, upper) and narrow-spiking SOM^+^ (NS-SOM^+^, Fig. 3A, lower right) cells, which occupied separate regions of the embedding. These sub-populations align with previous observations of SOM^+^ subtypes with specific synaptic targeting^45^ and differential behavioral functions^53,54^. Importantly, these SOM^+^ sub-populations would be difficult to identify using spike width alone, as SOM^+^ cells as a whole form a continuum on this metric (Fig. 3E, purple) that overlaps with both PV^+^ cells (Fig. 3E, teal) and numerous untagged cells, presumably including excitatory neurons (Fig. 3F). PhysMAP’s multimodal approach disambiguates these SOM^+^ subtypes while simultaneously distinguishing PV^+^ and putative excitatory cells.

This ability to optimally identify underlying cell types extracellularly also held for a separate heterogeneous dataset from CellExplorer^55^ spanning multiple brain areas. Here, the multimodal approach again separated cell types better than unimodal visualizations (Fig. S7A to E) and demonstrated superior performance from both supervised classification (Fig. S7F) and unsupervised clustering perspectives (Fig. S7G).

*These results establish that PhysMAP can identify cell types in extracellular recordings by effectively leveraging informative modalities while discounting less reliable ones*.

### PhysMAP identifies inhibitory cell types with accuracy greater than or equal to other unsupervised methods

Targeted optogenetic experiments like those in the previous section offer ideal conditions for cell type identification: during recordings, single electrodes are advanced until optically-responsive cells are found and isolated. However, in most electrophysiological recordings, electrodes take an unbiased sampling of all neurons. Typical recordings also capture neurons with poorer isolation, at variable distances to somata, and more variable light intensities than targeted tagging experiments. Cell identification in such settings faces additional challenges: opsin-expressing cells can show diverse responses including “paradoxical suppression”^56^, and non-expressing cells may show excitation due to polysynaptic effects. In order to demonstrate whether or not PhysMAP satisfies the fourth property of an ideal tool (identifying cell types in real use cases), we must first examine how it performs when optotagged cells aren’t specifically searched for and enriched within a dataset. We demonstrate that despite these challenges, PhysMAP outperforms single modalities and other unsupervised methods in identifying cell type structures without optogenetic targeting, while requiring less computation time and minimal architecture tuning.

We applied PhysMAP to a large dataset of optotagged inhibitory neurons collected using Neuropixels Ultra probes from mouse visual cortex^57^, containing waveforms and 3D-autocorrelograms (ACGs) from 8,953 putative single units including 237 PV^+^, 107 SOM^+^, and 118 VIP^+^ cells. PhysMAP’s embedding (Fig. 4A, top left) clearly separated PV^+^ cells (teal) from SOM^+^ cells (purple), while VIP^+^ cells (pink) were scattered among untagged presumptive excitatory cells (gray). In contrast, the 3D-ACG visualization showed poor clustering (Fig. 4A, top right) but waveform shape clustering was ostensibly comparable (Fig. 4A, bottom).

**Figure 4.**
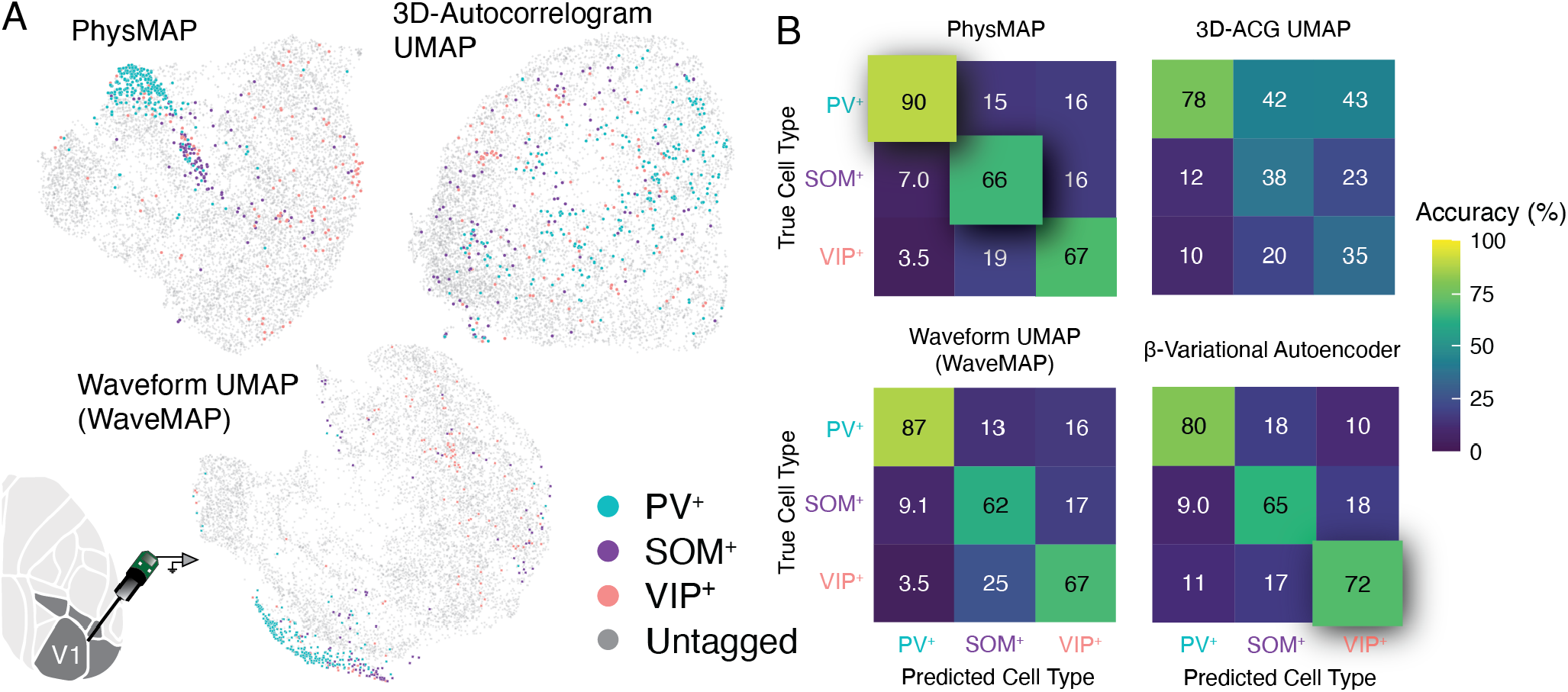
PhysMAP outperforms single-modality approaches and deep learning methods for inhibitory cell type identification. (**A**) UMAP visualizations of mouse V1 neurons recorded with Neuropixels Ultra probes, shown using different dimensionality reduction methods: PhysMAP multimodal integration (**top left**), UMAP on 3D-autocorrelograms alone (**top right**), and UMAP on waveforms alone (WaveMAP, **bottom**). Cells are color-coded by optogenetic tagging: PV^+^ (teal), SOM^+^ (purple), VIP^+^ (pink), and untagged cells (gray). **(B)** Confusion matrices comparing classification performance of a GBM (raw accuracy) trained on the high-dimensional representations (all matched at 60 dims.) across three approaches: PhysMAP (**top left**), 3D-ACG UMAP (**top right**), and waveform UMAP/WaveMAP (**bottom left**). A multi-layer perceptron was used to classify cell types from the latent embedding of a pair of *β*-variational autoencoders (**bottom right**). Values represent the percentage of cells from each true cell type (rows) assigned to each predicted type (columns). Color intensity corresponds to classification accuracy. Highest true positive percentages per cell type across all approaches are highlighted by embossing.

A GBM classifier trained on PhysMAP’s multimodal representation outperformed both unimodal representations of 3D-ACG and waveform shape across all three optotagged cell types (Fig. 4B). PhysMAP also outperformed classifiers trained directly on the concatenated raw modalities for most cell types (80%, 64%, and 71% for PV^+^, SOM^+^, and VIP^+^, respectively), demonstrating that its cell type-informative structure isn’t merely an artifact of optogenetic targeting.

We compared our approach against a *β*-variational autoencoder (VAE) method, representative of current unsupervised deep learning approaches. Although the VAE successfully reconstructed both waveform shapes and 3D-ACGs (Fig. S8A) and produced sensible embedding structures (Fig. S8B), a classifier trained on its embedding failed to outperform the GBM classifier trained on PhysMAP (Fig. 4B, bottom right). It is important to note that we used a single VAE architecture trained for each modality separately (Fig. S9), in contrast to extant ensemble approaches with multiple VAEs^30,58^, as the latter would be computationally infeasible to compare. Furthermore, although these alternative identification approaches are possibly very powerful (however, compare Fig. S10B, left to Supp. Fig. 6G of^30^), they are not ideal for the use case we intend for PhysMAP of being an iterable tool to quickly explore different datasets across several modalities: the ensemble approach would have entailed training ten separate models for each unit in the dataset. We find that PhysMAP is computationally more efficient. The *β*-VAE pair needed an expensive, high-performance GPU (NVIDIA L40S GPU; 500 epochs, 81 dimensions; 3.2 minutes) to achieve comparable training times to PhysMAP on a laptop (Apple M2 CPU; 500 epochs, 81 dimensions; 3.0 minutes) at inferior performance.

*Thus, PhysMAP provides superior multimodal representations with significantly less computational overhead compared to deep learning methods, making it ideally suited to data exploration and iteration across many modalities and datasets*.

### Multimodal visualization identifies batch effects

We’ve demonstrated that PhysMAP can uncover cell types within single datasets. For experimental utility, PhysMAP must consistently identify cell types across different animals and conditions. However, in real-world use cases, conditions differ between experiments, and it is possible that not all modalities maintain their usefulness due to batch effects. For example, when comparing anesthetized versus awake recordings, waveform shape may be conserved while spike timing modalities (such as ACG) differ significantly. Conversely, another dataset might use different signal processing filters within the same behavioral experiment, altering the relationship between waveform shape and cell type (see Fig. 5 of^42^) but preserving spike timing patterns. Such “distributional shifts” impede transfer learning when the relationship between cell type and properties changes as occurs across experiments. These shifts most commonly appear as lab-specific effects due to technical differences in data processing or experimental procedures^59^.

**Figure 5.**
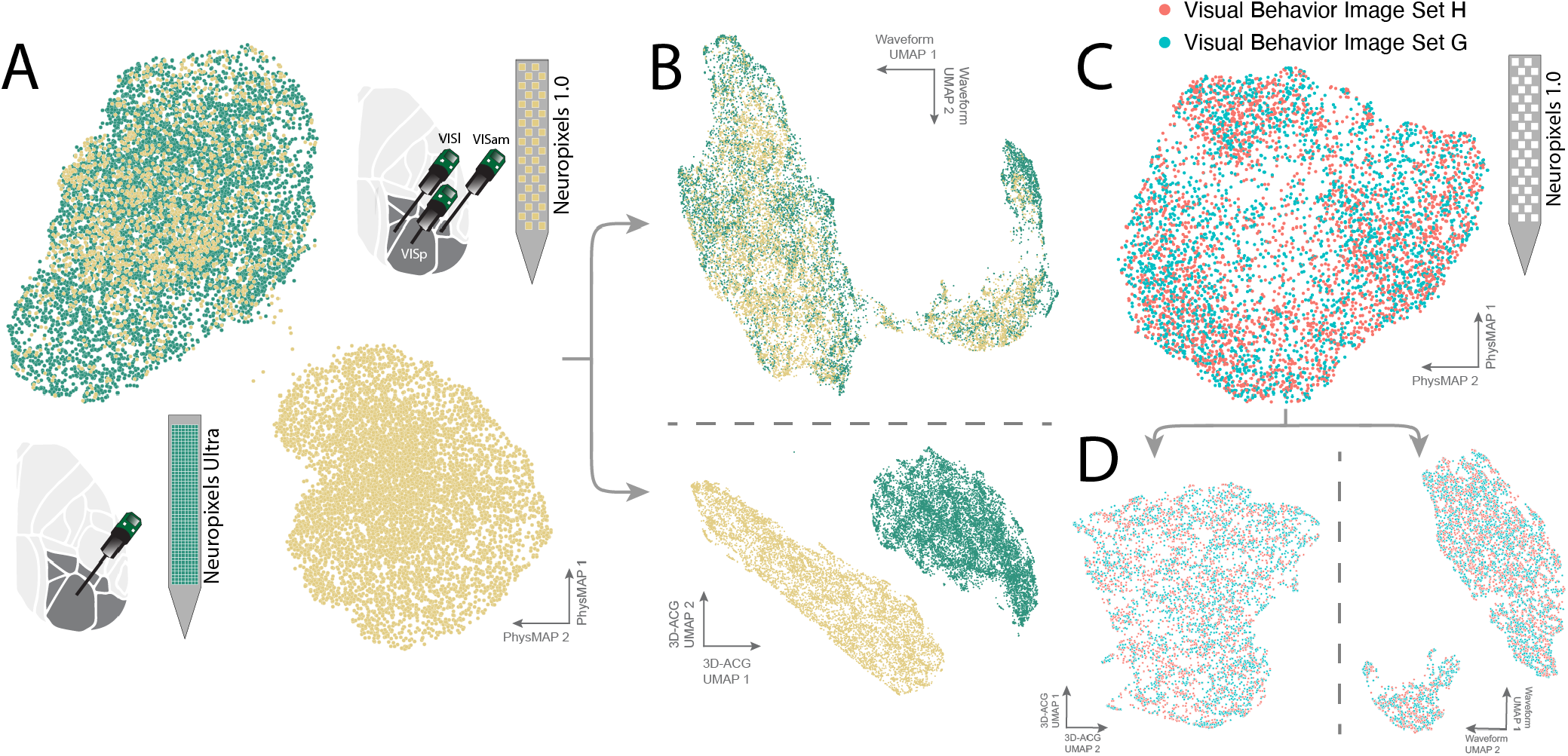
PhysMAP enables cross-dataset cell type identification through modality assessment and label transfer. (**A**) Two mouse visual cortex datasets visualized using PhysMAP: the Visual Behavior dataset (VB, green, recorded with Neuropixels 1.0) and the Ultras dataset (Ultras, yellow, recorded with Neuropixels Ultra). Diagrams show probe placements for each experiment. Waveform shape and 3D-autocorrelogam (ACG) are combined in PhysMAP and experiments are shown in the same visualization. (**B**) Constituent modality-specific UMAP projections with waveform shape (**top**) and 3D-ACG modality (**bottom**). Units are again colored according to originating experiment (green for Ultras; yellow for VB). Horizontal dashed line is just to make clear that top and bottom are part of different modalities. (**C**) Waveform shapes and 3D-autocorrelograms from two subsets (different animals) of VB experiments, one containing an image set H (in red) and another an image set G (in blue). (**D**) The VB multimodal representation is split into its constituent modalities. This consists of waveform shape (**left**) and 3D-ACG (**right**).

**Figure 6.**
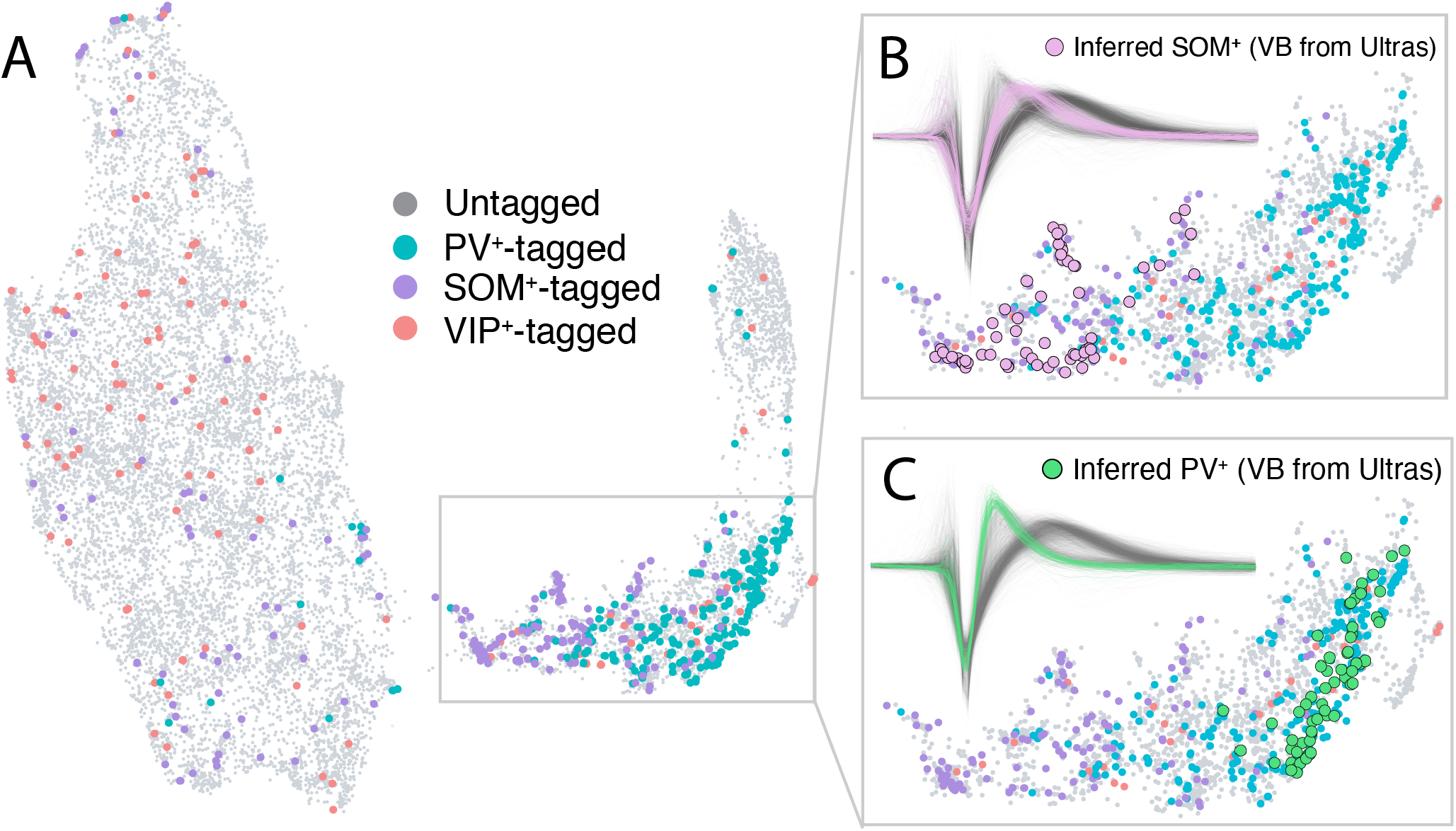
PhysMAP enables cross-dataset cell type identification through modality assessment and label transfer. (**A**) Cell type ground truth labels in the combined dataset embedding. Optotagged inhibitory interneurons are color-coded: PV^+^ (teal), SOM^+^ (purple), and VIP^+^ (red) cells, with untagged cells in gray. Inferred SOM^+^ cells in the VB dataset (pink circles) based on nearest-neighbor relationships with ground truth SOM^+^ cells from the Ultras dataset with waveform shapes shown (pink in inset) on a background of all waveforms in the VB dataset (gray in inset). (**C**) Inferred PV^+^ cells in the VB dataset (green circles) using the same approach also with waveform shapes shown (green in inset) on a background of all waveforms in the VB dataset (gray in inset).

These batch effects present a hidden danger wherein cell type identification is performed on a dataset in which the fundamental relationship between features and labels is different than in the dataset on which a classifier was trained. The danger increases with classifiers that simply output predictions without exposing underlying data distributional alignments. Thus, a key feature of any tool to identify cell types must be to detect batch effects and assess modalities for inclusion when performing cell type identification^60,61^. We experienced this deleterious effect firsthand with anchor alignment^62^ (used in a previous iteration of PhysMAP) which dramatically reduced classification accuracy by introducing random distributional shifts in noisy modalities (in this case, ACG; Fig. S10A). These effects reduced classifier performance by, on average, 22% across the three cell types investigated. This negative effect also appeared when PhysMAP was applied to cerebellar cell types (Fig. S10B;^30,63^). However, we do note that using PhysMAP without alignment precludes its application on new data without recomputing the WNN transformation. We show this approach does not incur any deleterious data leakage effects using the control analysis from^64^ (Appendix B).

To demonstrate PhysMAP’s ability to identify batch effects and assess modalities for inclusion, we examined two independently collected large-scale datasets: the previously investigated Ultras dataset and the Visual Behavior (VB) dataset^65^. To assess distributional alignment between these two datasets, we jointly projected them into PhysMAP’s multimodal embedding using normalized waveform shape and 3D-ACG (Fig. 5A), color-coding by dataset (Ultras in green, VB in yellow). While some units overlapped (gold units within the green cluster at upper left), most were completely separable—a batch effect made visible by PhysMAP’s unsupervised approach. Examining individual modalities (Fig. 5B), waveform shapes largely coincided between datasets (Fig. 5B, top), with minor differences: Ultras waveforms concentrated at borders relative to VB, and some Ultras units formed a separate cluster (middle right) composed of tri-phasic waveforms. These waveforms consisted of large pre-hyperpolarization peaks—likely waveforms from neurites that are visible only on very high-density probes^57^. In contrast, 3D-ACGs showed complete separation between datasets (Fig. 5B, bottom), likely because Ultras ACGs contained no spikes at certain timescales due to masking during optotagging stimulation. This indicates 3D-ACG should be discarded before further analysis. Thus, PhysMAP’s visualization-first approach represents a key advantage over purely classifier-based approaches that would have failed silently and led to erroneous conclusions within downstream analyses.

Note, while this dataset may have been reduced down to just one modality, we don’t mean to imply that any two experiments will not align on modalities such as 3D-ACG. The VB dataset is itself also divided into two experiments, each following the same protocol but using different natural image sets. These two datasets were indistinguishable both in the multimodal embedding space (Fig. 5C) and within each individual constituent modality (Fig. 5D). We emphasize that there are no extant datasets that we know of that have high-quality optotagging of all major inhibitory cell types during behavior, so while our analysis reduces to the unimodal case when examining the VB and Ultras datasets, it is an example of PhysMAP’s graceful handling of inappropriate modalities that would have instead failed silently if a purely classifier approach had been taken.

### PhysMAP can infer cell types using reference datasets

Guided by the previous section, we used waveform shape alone and visualized cells by optotagged identity. While most PV^+^ and SOM^+^ units occupied halves of one narrow-spiking cluster (Fig. 6A, lower right), numerous SOM^+^ and VIP^+^ cells were scattered throughout the larger cluster of wide-spiking excitatory cell types. The vast majority of neurons and cell types remain untagged since experimentalists can typically label only a few cell types, leaving most uncharacterized. This creates variable confidence across the neural population: regions with many tagged cells offer high classification confidence, while sparsely labeled regions remain uncharacterized (termed the “open-world problem” in machine learning^66^). And yet, in nearly all previous works on cell type identification, models are trained on datasets curated to only contain labeled cells. Cell type identifiers trained on datasets of only labeled cells fail to account for this experimental reality—they assign labels uniformly without considering out-of-distribution cell types or varying confidence based on local ground truth density. This observation highlights a critical gap between how most cell type classifiers are built and how they must function in practice. Can we develop an identification approach that acknowledges incomplete knowledge and selectively identifies cell types only where sufficient evidence exists? We answer this by creating a classifier that uses local densities of ground truth information exposed by PhysMAP.

We built a nearest-neighbor classifier based on the amount of ground truth labels surrounding every untagged neuron in PhysMAP’s representation. For each untagged VB unit, we calculated the proportion of nearest neighbors from the Ultras dataset belonging to each labeled cell type (including other untagged cells) on the multimodal graph constructed by PhysMAP (Fig. A.13J). This approach reduces the number of identified neurons but increases accuracy and avoids low-confidence regions (presumably composed of uncharacterized cell types). Based on this tradeoff (Fig. S11), we determined thresholds of 20% for nearest PV^+^ and 7.5% for nearest SOM^+^ neighbors, yielding 63 PV^+^ and 62 SOM^+^ units with 95% and 67% accuracy, respectively, all localizing to the narrow-spiking cluster shown by Fig. 6E and Fig. 6F, respectively. It should be noted that these classification accuracies are higher than in Fig. 4B because the selection of ground truth nearest neighbors acts as a “confidence threshold”^30^. As the proportion of ground truth neighbors increases, the accuracy of identification remains either roughly constant (SOM^+^) or monotonically increasing (PV^+^) except for when the number of untagged cells with a certain ground truth neighbor proportion reaches zero.

Examining the inferred SOM^+^ cells, we found they aggregated only in strongly SOM^+^-labeled areas and showed intermediate to narrow waveform widths matching ground truth SOM^+^ cells from previous analyses in this work. Similarly, inferred PV^+^ cells segregated to regions surrounded by Ultras PV^+^ cells and showed very narrow waveforms consistent with known PV^+^ properties. While we likely identified only a subset of these inhibitory neurons, we did so with higher confidence than possible from a simple classifier. PhysMAP’s visualization-first approach allows direct assessment of where and why units are classified, building trust through transparency rather than relying solely on blind predictions from classifiers trained on datasets potentially containing batch effects.

### Inferred types from PhysMAP match ground truth properties

In the previous section, we inferred PV^+^ and SOM^+^ cells by projecting VB and Ultras datasets into a shared multimodal representation and then using Ultras ground truth to predict cell types in VB. But do the properties of the inferred cells in VB match known properties of these cell types? To determine this, we examined the properties of these inferred PV^+^ and SOM^+^ cells.

In passive conditions, inferred PV^+^ cells exhibited fast firing rates, vigorous low-latency stimulus responses, firing rate deflections to the onset and offset of full-field flashes, and spike frequency spectra peaks in the gamma frequency band (Fig. 7A)—all known characteristics of these types of interneuron^31,32,67–69^. However, inferred SOM^+^ cells (Fig. 7B) displayed lower maximum firing rates, suppression to stimuli, and more variable responses than PV^+^ cells, matching previous findings^70,71^. Furthermore, their spike spectra showed peaks at lower frequencies consistent with their role in driving visually induced oscillations at lower LFP frequencies (< 30 Hz)^69^. These cell type-specific patterns were largely consistent across all cells of each inferred type, with PV^+^ cells showing strong responses to the onset and offset of stimuli (Fig. 7C) whereas SOM^+^ cells were more variable and sustained in response (Fig. 7D).

**Figure 7.**
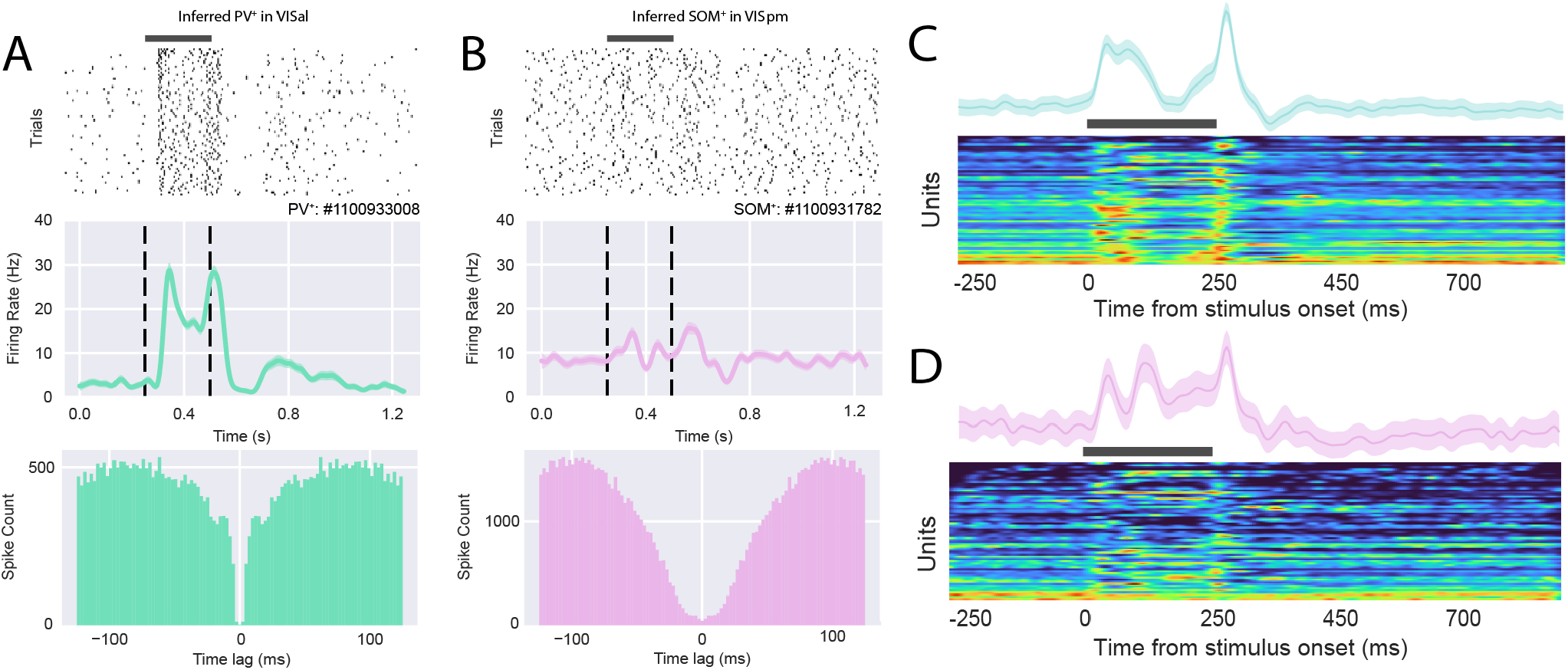
Label transfer allows inference of PV^+^ and SOM^+^ cell types. (**A**) Simultaneous raster plots in response to a dark full-field flash (**top**), smoothed trial-averaged firing rates (spikes/s with SEM and 100 ms time bin Gaussian smoothing; **middle**), and autocorrelograms (ACG; **bottom**) for simultaneously recorded inferred PV^+^ cell in visual areas VISal (green). Black overbar or vertical dashed lines indicate stimulus onset and offset times. Each raster’s row is a simultaneous trial shared by all cells. (**B**) Simultaneously inferred SOM^+^ cell in VISpm (pink, contemporaneously recorded with the PV^+^ neuron in Fig. 7A) also with rasters, smoothed trial-averaged firing rates, and ACGs (**top, middle**, and **bottom** respectively). (**C**) PSTH’s (normalized spikes/s with SEM) for all inferred PV^+^ units (across experiments). (**D**) PSTH’s for inferred SOM^+^ neurons. Each trace consists of binned spike times (1 ms, non-overlapping bins) and is normalized by the maximum trial-averaged firing rate for said unit.

Collectively, inferred PV^+^ and SOM^+^ cells match their expected physiological properties *in vivo*. These results provide evidence that PhysMAP can leverage ground truth datasets to enable the identification of cell types in datasets where ground truth is not available.

### Case study: Using PhysMAP to identify microcircuits in vivo

Currently, technical limitations mean that circuit dynamics are inferred by examining only single cell types at a time. However, neural computation is not the product of cell types operating in isolation but instead emerges from the interactions between different cells^72^. Having established that PhysMAP can identify several cell types simultaneously during behavior, we now demonstrate how this capability can be extended to the identification of microcircuits *in vivo*.

To demonstrate PhysMAP’s ability to identify microcircuits during behavior, we examined the relationships between PV^+^, SOM^+^ cells and directly optotagged VIP^+^ cells all simultaneously within the same experiment (Fig. 8A;^73,74^). Note that since VIP^+^ cells were not identifiable given the waveform shape modality alone (only with 40% accuracy [Fig. S11]), here they needed to be directly identified. As additional verification, the firing rate patterns produced by this VIP^+^ cell (Fig. 8A, top) are very similar to the PSTH’s of VIP^+^ cells documented elsewhere in the literature (compare Fig. 5G in^75^ to Fig. 8A, bottom).

**Figure 8.**
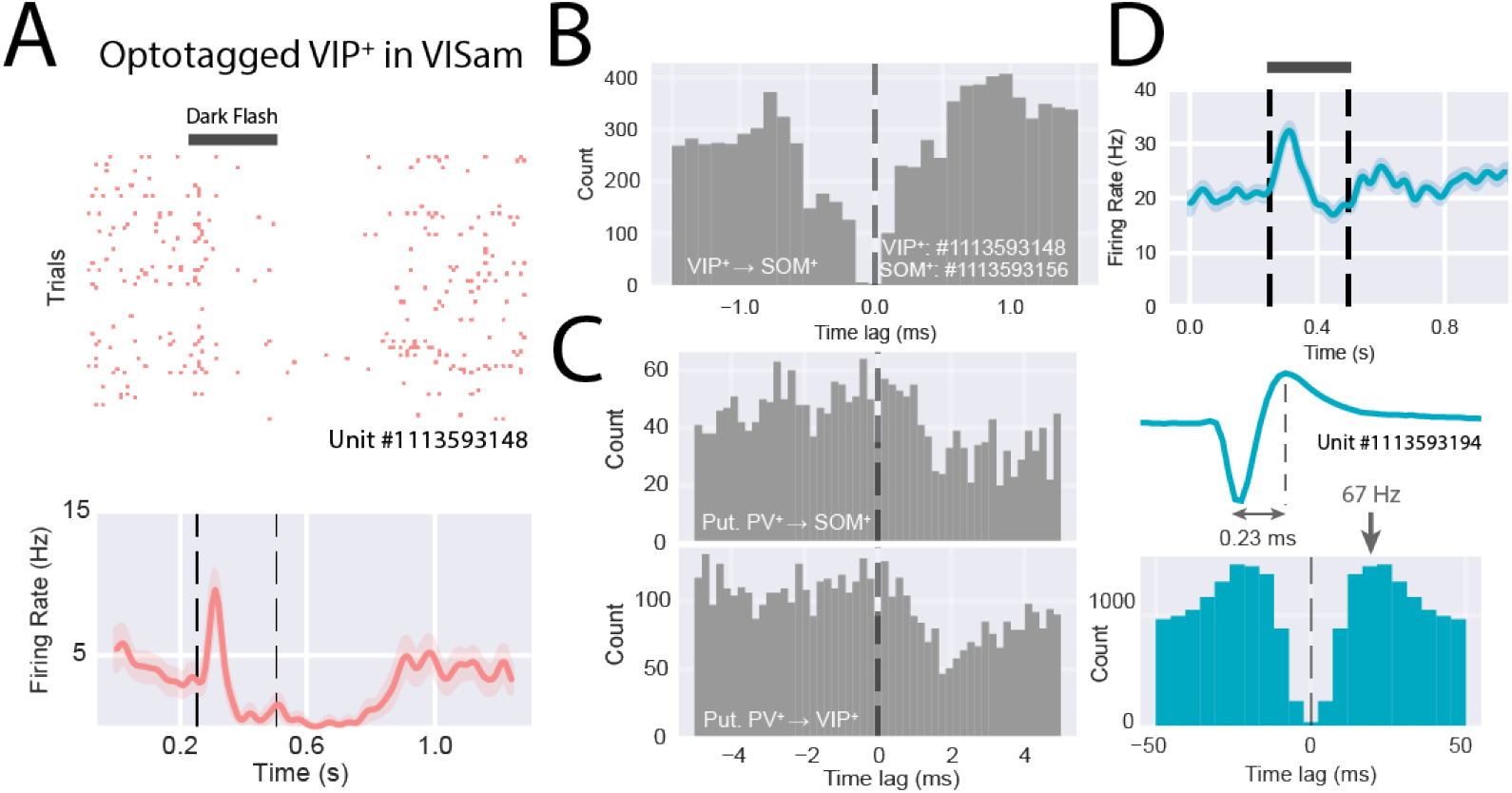
Simultaneous monitoring of three interneuron cell types with PhysMAP reveals microcircuits during behavior. (**A**) A simultaneously recorded optotagged VIP^+^ cell (red) during visual stimulus (black full-field flash) presentation with spike raster (**top**) and trial-averaged PSTH (mean with SEM; **bottom**). (**B, left**) Cross-correlogram with the VIP^+^ cell spike times (target) onto a SOM^+^ (second SOM^+^ cell in A) cell’s spike times (reference). Bins are 0.1 ms. Dashed line to indicate time bin zero. (**C, top**) Cross-correlograms for a putative PV^+^ cell onto the SOM^+^ cell in F (2 ms bin width). Dashed line indicates 0 ms bin. (**C, bottom**) Cross-correlograms for a putative PV^+^ cell onto the VIP^+^ cell in **F** (2 ms bin width). Dashed line indicates 0 ms lag. (**D, top**) Average spike waveform of the putative PV^+^ neuron inhibiting both the SOM^+^ and VIP^+^ cells. (**D, bottom**) 2D-autocorrelogram of the putative PV^+^ cell with peak in firing rate at 67 Hz (2 ms time bins).

Using pair-wise cross-correlations fitted with a generalized linear model of spiking interactions (GLMCC)^76^, we found that a SOM^+^ and VIP^+^ cell pair showed symmetric decreases in firing around time lag zero (Fig. 8B), indicating possible mutual inhibition (a ubiquitous motif)^75,77,78^ and/or simultaneous inhibition from an upstream interneuron. Investigating this latter hypothesis, we examined the cross-correlogram between a third neuron and the SOM^+^ and VIP^+^ cells and found that it suggested that the VIP^+^-SOM^+^ pair were strongly inhibited following this upstream cell’s spikes (Fig. 8C; top and bottom, respectively). The third neuron had fast firing properties and a vigorous response to stimuli Fig. 8D, top), a waveform with an extremely short trough-to-peak width (0.23 ms; Fig. 8D, middle), with a predominant peak in the autocorrelogram around 67 Hz (gamma band; Fig. 8D, bottom) strongly suggestive of this being a PV^+^ cell (although it was not identified by our algorithm).

These types of connections were not hard to find in high-density probe recordings. In this particular experiment, there were 78 putative monosynaptic connections (Fig. S12A) including both inhibitory and excitatory types (Fig. S12B, top and middle respectively), with some also being bidirectionally connected neurons mixing both inhibitory and excitatory connections (Fig. S12B, bottom). The VIP^+^ and SOM^+^ units made several inhibitory connections each, furthering our confidence that they have been positively identified as inhibitory types (units 6 and 7 in Fig. S12A).

These results suggest that not only can PhysMAP deal with batch effects and identify multiple untagged cell types, but it can also identify microcircuit interactions during behavior.

## Discussion

In this study, we developed PhysMAP, an approach that uses multimodal electrophysiological information to compute a representation for simultaneous cell type identification *in vivo*. PhysMAP optimally combines single modalities across seven datasets, outperforms comparable deep-learning methods with much less computation^30^, and enables analysis of the activity and interactions between multiple cell types simultaneously during behavior.

While pure classifier approaches might seem sufficient for cell type identification, three major barriers make them unsuitable: ubiquitous batch effects, scarcity of ground truth datasets, and uncertainty about which modalities to include. PhysMAP addresses all three prerequisites for trustworthy cell type identification.

First, PhysMAP visualizes dataset misalignments, providing a critical assessment of how appropriate a dataset is for cell type identification. This sanity-checking prevents erroneous conclusions from misapplied identification. Our experiments demonstrated dramatic accuracy decreases (by 22% and 36% in Fig. S10A and B respectively) when using datasets with misalignment due to batch effects. Direct identification does not make these effects visible.

Second, most electrophysiological recordings lack ground truth. Of the four known large-scale mouse experiments with optotagging, each covers only a single brain region. For primates and other relevant species, no such datasets exist. PhysMAP offers exploratory visualization of neuronal heterogeneity—for example, revealing SOM^+^ cell diversity across cortical areas, with both broad- and narrow-spiking waveforms as predicted by the literature^44,45,50^. Similarly, it revealed distinct properties of excitatory neurons across layers, aligning with observations from slice electrophysiological experiments^1^. This physiological variation within “discrete” cell types is invisible to many pure classifiers, but is readily apparent with PhysMAP.

Third, it is not well understood which modalities to use when building a reference dataset. PhysMAP offers an intuitive head-to-head comparison through visualization of each modality that pure classifier-based methods do not readily provide. For instance, in both juxtacellular and extracellular recordings, we show waveform shape is the most reliable modality for delineating ground truth cell type and ISI distribution, the least. These results reaffirm our previous work^24,25^ and are inconsistent with other work that emphasizes spike timing information. Moreover, PhysMAP provides this natively on a per-modality and per-unit basis, while interpretability in pure classifier methods necessitates retraining on data with modalities and units censored one by one.

Despite PhysMAP’s successes, our real-world application (Fig. 5) reveals important issues that need to be considered for cell type identification. Differences in tasks—even with similar probes in the same brain region— can produce pronounced batch effects in spike timing modalities. In addition, differences in signal processing filters during data acquisition can produce batch effects in waveform shape (as we’ve shown previously in^42^). We also recommend that ground truth generating experiments include standardized periods of “spontaneous activity” (involving no behavior) from which to generate spike timing modalities. These are easily replicable as they can be appended onto a lab’s existing behavioral workflow. Additionally, other behavior-independent modalities should be explored, such as spike-triggered LFP’s^58^ or spike-LFP coupling, which may separate PV^+^ and SOM^+^ types^79,80^. Other modalities might include specific “localizer” stimuli^24,58^ that generate distinctive activity patterns useful for identifying certain cell types. Other multimodal work has also shown that the multi-channel waveform “footprint” improves identification performance^81^. The multi-channel waveform was not investigated in this work as that would preclude cross-dataset cell type identification (different probe geometries between Neuropixels 1.0 and Ultras).

Our analysis revealed that few existing datasets have sufficient quality and comprehensive cell type coverage to serve as reference datasets. Only the Ultras dataset contained reliable tagging for all three major inhibitory types, and even this covers only the visual cortex of the mouse. High-quality single-unit recordings with optotagging remain challenging due to light attenuation, transgene expression specificity^82^., and low yield for rare cell types like VIP^+^ neurons which compose only ∼13% of inhibitory neurons^83^. Higher-density recordings^57^. and improved light delivery via waveguide optrodes^84^ may increase yields. Additionally, improved protocols for isolating directly tagged cells, such as using synaptic transmission blockers^30^, would further improve ground truth quality.

In addition, single unit isolation also remains challenging even with modern technologies^85^ and sorting methods^86^. A recent study^87^ predicts that on average, across 12 publicly available datasets, ∼24% of spikes from purported single units are false positives from nearby units. While this may not significantly impact population-level analyses^88^, it can blur physiological differences between cell types. For example, mixing fast-firing narrow-spiking PV^+^ cells with slow-firing broad-spiking excitatory cells can potentially create artifactual intermediate-property units, thus underscoring the importance of high-quality single unit isolation^24^. This mixing effect is evident when comparing unambiguous juxtacellular recordings (showing stronger cell type structure) with extracellular recordings (showing less structure) even in highly curated datasets.

Given these barriers presented, it might be tempting to ask why to choose to examine cell types with electrophysiological over optical approaches. Despite these challenges, electrophysiology offers several advantages over optical approaches for cell type characterization: simultaneous monitoring of multiple cell types on single trials, scalable access to cells across the brain, and exploration of cell types in primates and non-model organisms. While optical methods allow population monitoring, they currently limit imaging to only two fluorophores simultaneously^89^. Electrophysiology enables monitoring all cell types together, helping understand single-trial circuit dynamics^90,91^, and bridging anatomy and dynamics^92^. Optical methods also require optical access, limiting them primarily to cortical regions due to light attenuation in tissue^82^. Deeper imaging requires GRIN lenses or cortical aspiration, causing significant brain damage. Furthermore, optical methods are limited by their “photon budget”^93^: only a fixed number of photons are deliverable into the brain and so there exists a trade-off between number of units recorded and sampling rate^94^. Electrophysiology avoids these limitations, allowing arbitrary probe placement and simultaneous recording of thousands of neurons at high sampling rates^20^, with technological improvements promising even greater scaling.

Lastly, cell type-specific imaging requires transgene delivery and expression. PhysMAP only requires electrical activity recordings, making it valuable for studying non-human primates, where viral methods and optical access are extremely limited by technical constraints^13,14,95^. PhysMAP provides a low-risk solution for obtaining cell type information from routine electrophysiology in primates.

### Summary

PhysMAP offers robust exploration of putative cell types in unlabeled data and fast, efficient identification of cell types with ground truth. When used judiciously, it enables unsupervised identification of multiple cell types, in real-time, in behaving animals.

## Methods

The methods are organized as follows. We first describe the datasets that we used for the paper and then describe the weighted nearest neighbor method from PhysMAP. Next, we outline the various analyses used for quantifying the performance of PhysMAP relative to individual modalities and also other controls. Finally, we describe the label transfer method to identify cell types along with various analyses to verify their properties.

### Open Datasets

Here, we only briefly detail the most relevant aspects of each dataset for our purpose of validating our PhysMAP approach (Table 1). We refer the reader to each dataset’s respective publication for additional methodological information.

**Table 1:** The seven datasets analyzed with the modalities used and the cell types identified in each. Two datasets are contained in^96^

| Study | N | Used modalities | Labeled cell types | Brain areas | Data URL |
| --- | --- | --- | --- | --- | --- |
| Yu *et al.* 2019 | 246 | Waveform shape, ISI dist., PSTH | PV<sup>+</sup>-4, PV<sup>+</sup>-5, SOM<sup>+</sup>, E-4, E-5 | S1 | Link |
| Lakunina *et al.* 2020 | 373 | Waveform shape, ISI dist. | PV<sup>+</sup>, SOM<sup>+</sup>, E | S1 | Link |
| Petersen *et al.* 2020 (Allen Institute Visual Coding and Buzsaki Lab) | 430 | Waveform shape, ISI dist., ACG, spike metrics | PV<sup>+</sup>, SOM<sup>+</sup>, E, Axo-axonic (PV<sup>+</sup>), VGAT<sup>+</sup> (pan-inhib.), VIP<sup>+</sup> | Visual cortex and CA1 | Link |
| Ye and Shelton *et al.* 2024 | 8,953 | Waveform shape, 3D-ACG | PV<sup>+</sup>, SOM<sup>+</sup>, VIP<sup>+</sup> | Visual cortex | Link |
| AIBS: Visual Behavior | 5,582 | Waveform shape, 3D-ACG | SOM<sup>+</sup>, VIP<sup>+</sup> | Visual cortices | Link |
| Beau *et al.* 2025 | 3,301 | Waveform shape, 3D-ACG | PkC<sub>SS</sub>, PkC<sub>CS</sub>, MLI, MFB, GoC | Cerebellum | Link |

### Juxtacellular Mouse S1 Dataset

The juxtacellular dataset used in Fig. 1 and Fig. 2 and analyzed in section was collected by^32^ and downloaded from the associated file sharing portal (see Table 1). Recordings were performed in the primary somatosensory (barrel) cortex of Sst-IRES-Cre × Ai32, Pvalb-IRES-Cre/Pvalb-Cre × Ai32, or Vip-IRES-Cre × Ai32 mice. These experiments focused on *in vivo* juxtasomatic electrophysiological recordings using glass micropipettes. After the end of recording, the cells were filled with biocytin/neurobiotin. Filled neurons both acted as a verification of the recorded cell type via their morphology and also served as landmarks to align recording depths of each cell with cortical layers as determined by histology.

ChR2-expressing neurons were located during recording by observing laser-evoked spikes that occurred 1-2 ms after the onset of brief light pulses (5 ms) delivered at low frequency (5 Hz) for 3 out of every 10 s. Spikes were examined by eye and non-spike events were removed after projection in PCA-space. To control for drift, only waveforms between the 25th and 75th percentile of all spikes from a cell were used in further waveform shape analyses (average waveform and waveform shape metrics). Averaged waveform shapes and ISI distribution were extracted for each neuron from the *spkwaveform_all*.*mat* file in the corresponding cell type structs. In addition, the waveform shape metrics of spike width and peak-to-trough ratio were also collected from this file. Cells labeled “fake SOM^+^” were reassigned to the PV^+^population and cells labeled “putative SOM^+^” were reassigned to SOM^+^. This reassignment was based on a careful investigation in^32^ confirming earlier findings that off-target recombination occurs in fast-spiking neurons of somatostatin-IRES-Cre mice possibly due to transient Cre expression during development^97,98^. PSTHs were calculated only from whisker touches during active whisking and activity resulting from both whisker protraction and retraction were used. Spiking was aligned to touch-onset and a PSTH calculated using 1 ms time bins. As not all cells contained PSTHs, only the subset of units containing all three modalities (waveform shape, ISI distribution, and PSTH) were used. To create the “concatenated” modality, we simply concatenated the waveform shape, ISI distribution, and PSTH for a given unit into a single long feature vector. Waveform metrics—peak-to-trough duration and peak-to-trough ratio—were calculated in the originating publication as the time from the first peak to the first trough and the ratio of absolute values of the peak to the trough respectively.

### Extracellular Mouse A1 Dataset

The extracellular data we analyzed from^52^ was downloaded from GitHub (see Table 1). Recordings were collected via silicon probes (A4×2-tet configuration from NeuroNexus) and light was delivered via optic fiber at around 200 *µ*m from the first recording site. PV^+^and SOM^+^ cells were identified by optotagging in Pvalb-Cre or Sst-Cre mice respectively that were the progeny of crossing LSL-ChR2 mice. Photoidentification of ChR2-expressing cells was relatively conservative with positively identified cells being those that exhibited significant firing rate changes (p < 0.001) in firing rate during the first 10 ms of stimulation-onset. Putative excitatory units were identified by a significant decrease in firing rate during the experiments in Sst-Cre mice. Recordings were collected from primary auditory cortex via silicon probe and stimulation was via an optical fiber located at the top of the probe recording sites. Spikes were sorted offline in Klustakwik and the units used in further analyses only if ISI violation rate (number of spike intervals < 2 ms) was less than 2% per cluster. In the original publication, only cells with spike quality index (SQI; ratio between the peak amplitude of the waveform and the average variance, calculated using the channel with the largest amplitude) above 2.5 were used; in our analysis, we only used cells with SQI above 4.0. Mean waveforms were calculated for a unit using the channel with greatest amplitude of spikes.

### Extracellular Mouse Visual Cortex and Hippocampus Dataset (CellExplorer)

The third dataset was obtained from the CellExplorer package^96^ and is heterogeneous, being composed of two separate extracellular electrophysiological data sets from the Allen Institute’s Brain Observatory and the Buzsaki lab, and included waveform shape, ISI distribution, autocorrelogram (ACG), and various derived electrophysiological metrics. We used the derived eletrophysiological metrics that all units shared and this included the following eleven:

- Spike width (*troughToPeak*)
- Peak-to-trough derivative (*troughtoPeakDerivative*)
- Pre-hyperpolarization peak-to-post depolarization peak ratio (*ab_ratio*)
- Coefficient of variation (*cv2*) measuring ISI distribution
- ACG tau rise (*acg_tau_rise*)
- ACG tau decay (*acg_tau_decay*)
- ACG tau bursts (*acg_tau_burst*)
- ACG refractory period (*acg_refrac*)
- ACG decay amplitude (*acg_c*)
- ACG rise amplitude (*acg_d*)
- ACG burst amplitude (*acg_h*).

More information on these metrics are available on the CellExplorer website docs (link). For the analyses, these metrics were concatenated for each neuron into a single vector per-unit to form the electrophysiological metrics modality. Given the large number of publications and protocols that have contributed to the CellExplorer dataset, we only provide a brief overview here.

### PhysMAP Preprocessing

Modalities from all neurons were preprocessed via the same steps required by and performed using Seurat v4:

1. **PCA reduction**: All modalities were initially dimensionality reduced to the smallest input modality dimension since all modalities into Seurat must be the same dimensionality. For the S1 dataset, this is 30 dimensions; for the A1 dataset, this is 20 dimensions; and for Ultras and Visual Behavior, this is 81 dimensions.
2. **Normalization**: Each modality is separately normalized using a centered log ratio transform.
3. **Rescaling and centering**: Each feature of each modality is linearly rescaled to occupy the same unit variance. Each feature is then centered by mean subtraction.

### Weighted Nearest Neighbor (WNN) Algorithm

The PhysMAP approach uses the WNN algorithm available from Seurat v4^38^ which is summarized as the following steps.

1. **Nearest neighbor identification**: Within the preprocessed, high-dimensional space of a given modality (here, waveform shape Fig. A.13A), a single unit is selected (Fig. A.13B, blue sphere) and its nearest neighbors identified via Euclidean distance in the ambient (un-dimensionality reduced) space (Fig. A.13B, red spheres).
2. **Nearest neighbor prediction (within modality)**: A prediction of the selected single unit’s waveform is made via unweighted averaging the *k*-nearest neighbors identified in Fig. A.13B. By default, we use *k* = 20 and Euclidean distances in the ambient space to locate the nearest neighbors.
3. **Within-modal affinity calculation**: The unit’s nearest neighbor predicted waveform shape generated by *k*-nearest neighbors 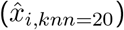 and its actual waveform shape (*x*_*i*_) are then passed into a modified UMAP distance kernel,

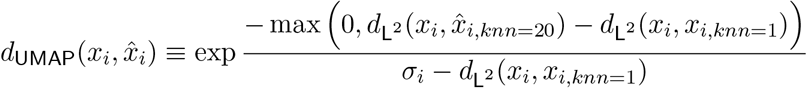

where 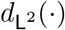 is the Euclidean (L^2^-norm) distance metric; 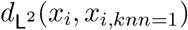 is the Euclidean distance from *x*_*i*_ to its nearest neighbor; and *σ*_*i*_ is a modified UMAP bandwidth equal to the average of the Euclidean distances from the *i*^th^ unit to the 20 nearest units with lowest non-zero Jaccard index. This equation is used to calculate the “within-modal affinity” which gives a measure of how predictive a certain modality is of this unit. This forms the numerator of the “affinity ratio” shown in Fig. A.13D.
4. **Nearest neighbor identification (cross modality)**: Examining normalized single unit ISI distributions (the “cross-modal” space) generated from the same neurons in Fig. A.13E, we locate the same unit and nearest neighbors previously identified but in ISI distribution-space Fig. A.13F.
5. **Nearest neighbor prediction (cross modality)**: As in Fig. A.13C, a nearest neighbor prediction is made but this time, in a different modality albeit using the same neurons identified in the original modality Fig. A.13G, red dashed circle.
6. **Affinity ratio calculation**: This unit and its prediction in the cross modality (ISI distribution-space) are used to compute the “cross-modal affinity” by passing it into a modified UMAP distance kernel; this forms the denominator of the waveform affinity ratio for a given unit (*S*_Wave_(*i*)) shown in Fig. A.13D. This process is repeated for every unit in the dataset and also in the reverse manner: beginning with the ISI distribution-space as the within modality and with the waveform shape-space as the cross modality. For datasets with more than two modalities, the affinity ratio is calculated for each modality versus every other and for every unit. Thus for a dataset with *d* data points and *n* feature modalities, there will be 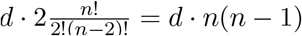 affinity ratios to calculate.
7. **Converting affinity ratios to modality weights**: With affinity ratios calculated for both waveform and ISI distribution across all units, modality weights (*β*_Wave_(*i*) and *β*_ISI_(*i*)) for each unit can be calculated. For a given unit, this weight is the ratio of a given modality’s affinity ratio for that unit divided by the sum of all other modality’s affinity ratios for that unit. For the two-modality case, this is summarized by the equation in Fig. A.13H.
8. **Calculating unit pair-wise affinities**: For every pair of units, an affinity is also calculated by passing both of them and their coordinates into UMAP’s distance kernel. This occurs for the same pair of units in both the waveform shape-space Fig. A.13I, top and the ISI distribution-space Fig. A.13I, bottom to determine pair-wise affinities *θ*_Wave_(*i, j*) and *θ*_ISI_(*i, j*) respectively. This is done for all pairs of points and all modalities.
9. **Creating the WNN**: To create the weighted-nearest neighbors representation, a modality-weighted sum of the pair-wise affinities Fig. A.13J, top is taken to produce a “connectivity matrix” Fig. A.13J, bottom. The *k*-nearest neighbor algorithm is applied to this matrix to form the final WNN (default number of neighbors is *k* = 200) which is then visualized into two dimensions with UMAP’s force-directed graph layout projection for visualization.

### Leiden Community Detection

To calculate the MARI score for each dataset across different numbers of clusters we used Leiden clustering^48^ on each dataset’s WNN graphs with resolution between 0.1 and 3.0 in 0.1 resolution intervals. The Leiden algorithm is a method for “community detection” which finds highly inter-connected nodes on a network graph akin to clustering in a metric space. Algorithms like Leiden (and the simpler Louvain algorithm) attempt to find a partitioning of the network into a set of communities that maximizes the modularity of each community. This modularity, ℋ, is a measure of how inter-connected the nodes are within a community versus outside of it. In the below definition, *m* is the total number of edges in the network, *e*_*c*_ is the number of edges in a community *c, γ* is the resolution parameter, and *K*_*c*_ is the sum of the degrees of the nodes in community *c*.

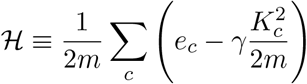

Thus, the resolution parameter is effectively a prior on the number of expected communities that should be found during the optimization; with lower resolution, less communities are found and with higher resolution, more communities are found.

### Modified Adjusted Rand Index (MARI) Calculation

Once this partitioning of a graph into communities is computed, a MARI score^49^ is calculated between each of these each neuron’s community membership and the ground truth cluster identity to determine how closely they corresponded. To understand MARI, which is a modification of the adjusted Rand index (ARI), we begin with an explanation of the Rand index. Given two clusterings, *X* and *Y* of the same dataset, the number of pair-wise elements that share cluster membership in both *X* and *Y* is *a* and the number of pair-wise elements that share different cluster memberships is *b*. The Rand index (*RI*) is 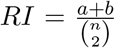 where *n* is the total number of pairs. The Rand index can also be interpreted as the sum of the number of true positive and true negative correspondences divided by the total number of guesses if one clustering is regarded as a classifier prediction of the other. However, in the classifier interpretation, the Rand index is not corrected for chance predictions. The ARI corrects for these chance predictions by subtracting the expected Rand index under the hypergeometric distribution given two independent clusterings and fixed number of clusters in each. MARI further refines this metric by instead incorporating a multinomial distribution which does not enforce cluster size and is a better assumption given that most clustering algorithms (including Leiden) do not fix cluster sizes. This was used to compare each of the Leiden clusterings at different resolutions to underlying ground truth clusters.

### Peri-stimulus Time Histogram Calculation

For each Leiden cluster in Fig. 2D, a peri-stimulus time histogram (PSTH) was calculated by aligning times to each whisker deflection onset and binning spikes into 10 ms bins. This was then normalized by subtracting the baseline spike rate for each neuron and then dividing by the standard deviation of each unit’s PSTH. These were then smoothed by a 5-point moving average and then grouped according to Leiden cluster membership for between -10 and 50 ms before and after the whisker deflection.

### Identification Algorithm

After a WNN is constructed for a dataset, we used UMAP’s force-directed graph layout procedure to project the graph into a high-dimensional embedded space. Dimensionality for this embedding was maximized to provide the greatest amount of information available for classifier comparison (the WNN algorithm only allows for a maximal output dimensionality equal to the highest dimensional input modality). We then used a random 15-85% test-train split and trained a stochastic gradient boosted tree model (GBM) classifier on the training set with five-fold cross-validation using the *caret* package in R^99^ to identify each underlying cell type with a multiclass (one-vs-rest) objective function. In some datasets, cell type classes with less than 10 examples were excluded because this would not guarantee that a cell would appear in both the training and test sets at the test-train split used. We then repeated this train-test split 20 times and averaged the balanced accuracy over these independent runs. Each classifier also underwent hyperparameter tuning via a grid search. This procedure was run identically for the controls in Fig. S5 except with varying graph embedding dimensions and using different classifiers available in the *caret* package. This same approach was used for the three modality case in Fig. S4C.

### Gaussian mixture model analysis of spike metrics

A Gaussian mixture model Fig. S2 was fit to units in^32^ by applying the *mclust* library’s *Mclust* function to the log-scaled peak-to-trough and log-scaled spike width metrics for each unit as done previously in the literature^23,25^. This function simultaneously compares many GMM models and finds the optimal number of clusters for each via the Bayesian information criterion (BIC). It arrived at a VVE (ellipsoidal with equal orientation) model and three clusters.

### β-variational autoencoder and multi-layer perceptron identification architectures

Using the Ultras dataset, a *β*-variational autoencoder (VAE) was constructed using a standardized architecture for each modality Fig. S9: a first/second 1-D convolutional layer (*N* = 64, *C*_in_ = 1*/*32, *C*_out_ = 32*/*64, *K*_size_ = 3) followed each by ReLU non-linearities and flattened; passed into an 81-dimensional latent embedding parametrized by mean *µ* and log variance log *σ*^2^ (linear transform layers, latent dim. = 81, hidden dim. = 64); then unflattened and passed into a decoder composed of a first/second 1-D convolutional layer (*N* = 64, *C*_in_ = 64*/*32, *C*_out_ = 32*/*1, *K*_size_ = 3) with a ReLU in-between. Networks were trained with the Adam optimizer over 200 epochs, grid search optimized for *β* based upon classification accuracy, and with logistic Kullback-Liebler (KL) divergence annealing. The *β*-VAE was optimized using the evidence lower bound objective function (ELBO) balancing both *l*^2^ norm reconstruction error and KL-divergence multiplied by the tunable *β* parameter. For the waveform shape VAE, *β* = 0.05 and for the 3D-ACG VAE, *β* = 0.1; notice that these optimal *β* reduce the VAE to nearly a vanilla autoencoder. The multi-layer perceptron (MLP) classifier (scikit-learn’s *MLPClassifier*) was composed of an input layer (10 units), hidden layer (100 units), ReLU non-linearity, final input layer (4 units), and softmax output for cell type prediction. The MLP classifier was trained for 1000 iterations with early stopping.

### Identification comparison

For each confusion matrix in Fig. 4B, the projection and identification process was trained 25 times for each of PhysMAP, waveform shape, 3D-ACG, and the *β*-VAE. For each iterate, the transformation was applied to the entire dataset and then a seed selected to determine a random 90-10 train-test data split. Across ten random splittings, five-fold repeated cross-validation was applied and averaged (across repeated CV’s and splittings) to determine each cell type’s raw classification accuracy. We also followed this same procedure for the confusion matrices in Fig. S10 with and without erroneous alignment. This procedure was also applied to PhysMAP with and without alignment on the C4 dataset shown in Fig. S10B^63^. The alignment procedure is not used in further iterations of PhysMAP but is documented for comparison and to demonstrate the deleterious effects of misaligned data distributions. The alignment procedure is explained in the next section for clarity.

### Reference mapping with anchors and cell type identification

Unlike the previous identification analyses, an 80-20 train-test split was conducted *before* applying PhysMAP. This was done to combat bias in cross-validation because of data leakage due to pre-processing transformations^64^. The training set was used to create the reference mapping upon, and the test set was used to create the query dataset. To construct the reference mapping, a WNN was constructed from each neuron’s waveform shape, ISI distribution, autocorrelogram, and derived electrophysiological metrics as before. Following the procedure for multimodal reference mapping recommended in Hao & Hao et al. 2021, we first ran supervised principal component analysis (SPCA) on the reference data. Next, the query dataset is projected onto the reference using the previously computed SPCA transform. Now that reference and query datasets occupy the same space, “anchors” can be calculated between the reference and query. These anchors are pairs of cells between the reference and query that are located within each other’s neighborhoods; this concept is also referred to as mutual nearest neighbors (MNN). These neighborhoods are defined by computing *k*-nearest neighbors with *k* = 5. This nearest neighbor search is conducted in only the top-30 SPC’s. Once all anchors are found, they are scored, which is an assessment of how confident we are in their correspondences. This score is calculated for each anchor given the 30 nearest neighbors in each reference and query dataset for both the reference and query data points of the anchor, respectively. The overlap of these nearest neighbor matrices between the 0.01 and 0.90 quantiles is then linearly rescaled to be between 0 and 1 for all reference-query anchor pairs; this provides a score for each anchor. A weight matrix is then constructed between each query cell *c* and each *i*th nearest anchor *a*_*i*_ (where *i* ∈ [1, 50]) based upon the distance to each query cell and the corresponding anchor score. These weighted distances *D*_*c,i*_ are calculated as

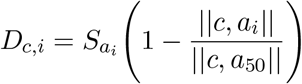

where 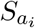 is the score of the *i*th anchor where || · || is the Euclidean distance. These distances are passed through a Gaussian distance kernel 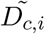 as,

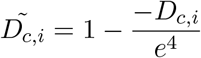

to form the entries of a weight matrix

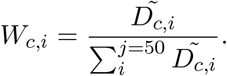

For cell type identification, a classification matrix *L* is created where each row corresponds to a ground truth cell type class and each column corresponds to each reference anchor. If a certain reference anchor pertains to a certain cell type, it is given a 1 and a 0 if otherwise. Label predictions *P* are then computed simply as,

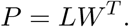

This returns a prediction score for each cell in the query dataset. A query cell is thus given a predicted cell type of whichever class has the highest prediction score.

### Pre-processing of the Visual Behavior dataset

The Visual Behavior dataset, containing the Images H and Images G experiments, was downloaded using the Allen Institute’s *AllenSDK* package. Units were selected only if they were from primary visual cortex and higher visual areas (HVAs). Furthermore, we were quite stringent with quality control: units were only selected for analysis if they had a signal-to-noise ratio (maximum amplitude of the mean spike waveform divided by the standard deviation of background noise) exceeding 3; a maximum of inter-spike interval violation over firing rate ratio of 0.05 (to limit the false discovery rate;^87^); a minimum presence ratio of 0.98 (the proportion of time a unit was present in a recording); and a baseline firing rate of 2 spikes\s over the entire recording. In addition, optotagged units were selected if their firing rose to twice their baseline during the first 10 ms of laser onset and were driven to a firing rate of at least 25 spikes\s.

The 3D-ACGs were calculated by taking the spike times across the entire experiment except for the optotagging epoch which occurred at the end of the session. The 3D-ACGs used a 1000 ms window size and a 10 ms bin size and were log-transformed (matching the Ultras dataset) and were constructed using the *NeuroPyxels* function, *fast_acg3d*^63^.

### Joint embedding and label transfer of Ultras and Visual Behavior

Both Ultras’ and Visual Behavior’s waveform shape and 3D-ACG modalities were concatenated and passed into PhysMAP. The multimodal representation was generated using *caret* with centered log ratio transform normalization on a per-unit basis. UMAP was calculated for 20 components, but the 2 component projection is used for the visualizations shown in Fig. 5A. The same approach was used for the individual modalities shown in Fig. 5B, top and bottom. For the multi-dataset nonlinear embedding of waveform shape (unimodal PhysMAP), shown are the optotagged cell type identities (PV^+^, SOM^+^, and VIP^+^) from each experiment amidst a background of untagged neurons Fig. 6E. Labels for Visual Behavior were generated using the previously described procedure (Methods: Pre-processing of the Visual Behavior dataset); labels for the Ultras dataset were generated by applying density-based spatial clustering of applications with noise (DBSCAN) to a UMAP embedding of PSTH responses in the first 10 ms after laser-onset^57^.

For label transfer from Ultras tags to Visual Behavior, we used a nearest neighbor approach: for a fixed number of neighbors (k = 33) for each Visual Behavior unit within the high-dimensional graph constructed within PhysMAP, we calculated what percentage of neighbors (including unlabeled cells) contains a given label from the Ultras dataset and set a minimum percentage threshold belonging to a unitary cell type label (20% for PV^+^ cells and 7.5% for SOM^+^ cells) in order for a label to transfer. These inferred cell types are shown in (Fig. 6F and G). This percentage can be adjusted to the desired level of confidence; accuracy increases with an increasing percentage of neighbors in the Ultras dataset with accuracy measured as the number of correct identifications if only the Ultras dataset is considered Fig. S11. The procedure was fixed at *k* = 33 and repeated 25 times for each percentile.

This method worked well for both PV^+^ and SOM^+^ cells, but because VIP^+^ cells were only very sparsely labeled and seemingly non-differentiated from broad-spiking excitatory cells, we used the direct labels obtained during the Visual Behavior experiments. Furthermore, VIP^+^ cells were poorly identifiable even at the highest confidence thresholds (a maximum accuracy of 40% at 8% nearest neighbor threshold; Fig. S11, red lines).

### Firing rate analysis of inferred cell types

For each peri-stimulus time histogram in Fig. 8A, bottom, firing rates were determined by collecting spikes into 100 ms time bins, summed across trials, and then smoothed by a Gaussian kernel with a standard deviation of the number of total bins divided by 6. The same procedure was done for each individual inferred unit in the Visual Behavior dataset and normalized (dividing by the maximum firing rate) to generate Fig. 8B.The stimuli used were a gray to black flash on a computer monitor for 100 ms across most of the visual field contralateral to the recorded hemisphere. In Fig. 8D, a natural image was used as the stimulus aligned instead. This image was one that was correctly identified as a change from the previously shown natural images indicated by the mouse licking a lickspout during the reward window.

For the autocorrelograms in Fig. 8D, bottom, spikes were gathered into 5 ms time bins and shown for a window 150 ms before and after 0 lag. To generate the trial-averaged spike spectra in Fig. 8E, the procedure in the chapter titled “Analysis of Rhythmic Spike Train Data” from^100^ was used. Specifically, this is a multitaper spectral density estimation method using 4 Slepian sequences (*scipy*.*signal*). The “pre-stimulus period” was taken as the 350 ms before to 100 ms after stimulus onset and the “post-stimulus period” is taken as the 100 ms after to 550 ms after stimulus onset. The reason for not aligning the pre- and post-stimulus periods directly with stimulus onset is the approximately 100 ms propagation time of stimulus information from retina to visual cortex (intuitively seen as a change in activity across most cell types). These were divided into both “hit” and “miss” trials Fig. 8E, middle and bottom in which the animal correctly identified the natural image as different from the ones before or else failed to respond to the changing image (indicated by a lick or lack thereof) respectively.

To generate the CCGs in Fig. 8F and G, we used a window/bin size of 3 ms/0.1 ms for the SOM^+^ to VIP^+^ CCG and 10 ms/0.2 ms for the VIP^+^ and SOM^+^ onto putative PV^+^ respectively. The ACG for the putative PV^+^ cell is shown with 100 ms window size and 5 ms bin sizes.

### Identifying putative monosynapses with a GLMCC

To identify putative monosynaptic connections between neurons, we examined each pair of neurons for low latency responses on the timescale of synaptic transmission delays. We used the revised general linear model on cross-correlogram (GLMCC) approach by^76^ which tends to perform better than both using classical cross-correlogram analysis^101^ and jittered cross-correlations^102^ in terms of minimizing both false positives and false negative rates. Furthermore, the GLMCC also yields an estimate of the strength of synaptic connectivity other than the classical and jittered methods, which only yield a binary identification or non-identification of a monosynaptic connection. More specifically, the underlying spike rate of a given neuron over time *c*(*t*) is given as the product of several exponential distributions modeling respectively slow time-varying fluctuations affecting each neuron in a pair *a*(*t*), the connection from the first neuron to the second *J*_*ij*_, and the connection of the second neuron to the first *J*_*ji*_.

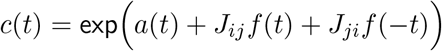

Here, *f* (*t*) is modeled as,

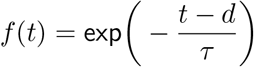

where *d* is the synaptic delay and *τ* is the synaptic timescale; the former is set to be between 1 and 4 ms while the latter is set to be 4 ms. Note that the underlying spike rate *c*(*t*) is modeled as a Poisson point process such that the spike probability *p*(*{t*_*k*_*}*|*θ*) over time with,

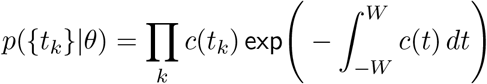

with a parametrization of *θ* = {*a*(*t*), *J*_*ij*_, *J*_*ji*_} and *W* being the integration window (set to 50 ms in our analysis). The prior distribution *p*(*θ*) is proportional to the following,

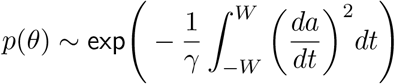

with the parameter *γ* setting the “flatness” of the *a*(*t*) is set to 2×10^−4^ in our analysis. The posterior distribution *p*(*θ*|*{t*_*k*_} was calculated given Bayes’ theorem and a maximum *a priori* (MAP) procedure used to calculate the parameters *θ*.

A monosynaptic connection is said to exist if the null hypothesis is rejected; the null hypothesis is that two neurons generate spikes at their baseline firing rate independently of each other. The variance of a Poisson process within a time interval Δ is equal to its mean and thus the mean number of spikes *n* is *n* = *c*(0)Δ. If *J* is small then the average number of spikes caused by a connection over an interval Δ is,

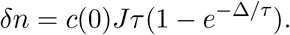

The condition that a monosynaptic connection is found is given by 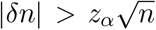 where z_α_ = 3.891 for the *α* = 0.0001 used in this analysis. Isolating the estimated connection strength 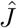 by substituting 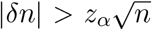 into the previous equation yields,

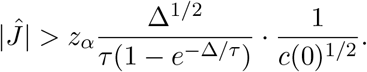

The variable Δ reaches its minimum at Δ = 1.26*τ* which reduces the previous equation to the following,

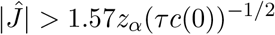

for which a connection is determined to be monosynaptic.

## Data Availability Statement

Post-processed data used in analyses and figure generation are provided in the repository https://github.com/EricKenjiLee/PhysMAP_Manuscript. Raw datasets are available from each source publication’s open data repositories in Table 1.

## Code Availability Statement

MATLAB, Python, and R for open dataset analysis and figure generation are available at https://github.com/EricKenjiLee/PhysMAP_Manuscript.

## Author Contributions

EKL and CC jointly developed the idea of a cell type classifier and ultimately a lookup table based on ideas from Dr. Karel Svoboda, and Dr. Josh Siegle. AG helped with derived waveform metric analysis and several control analyses. GH helped collect one of the open datasets and provided technical advice, especially with regards to the usage of quality metrics, and helped edit initial manuscript drafts. HY and CH provided many useful discussions, suggested analyses, and provided access to several datasets. AL and SJ provided technical support and also collected the S1 open datasets. AS, SO, and NS provided technical support and collected the Ultras dataset. PFP provided guidance on various approaches for multi-modal integration and other technical advice. EKL curated datasets. CC wrote initial R code and performed some analyses. Code was further refined and augmented by EKL. Additional analyses also written by EKL. EKL made figures and wrote initial drafts of the paper with CC. All authors edited the manuscript and provided feedback on presentation and clarity.

## Acknowledgments

We are grateful to Dr. Karel Svoboda, Dr. Anna Lakunina, and Dr. Josh Siegle for the initial ideas of cell type classifiers for electrophysiology. We also thank Dr. Adam Smoulder, Dr. Yujin Han, Dr. Josh Siegle, Pierre Boucher, Nicole Carr, Tian Wang, Vivan Moosmann, Tushar Arora, Mateo Umaguing, and Munib Hasnain for their thoughtful comments on our manuscript.

CC was supported by an NIH NINDS R00NS092972, R01NS121409, and R01NS122969 award; the Moorman-Simon Interdisciplinary Career Development Professorship from Boston University; the Whitehall Foundation (2019-12-77); and the Young Investigator Award from the Brain and Behavior Research Foundation (27923). The auditory cortex dataset (collected by AL and SJ) was supported by an NIH NIDCD R01DC01553. SJ was also supported by an NIH NINDS RF1NS131993. EKL was supported by an NIH NINDS F31NS131018. The Neuropixels Ultras dataset was supported by NIH NINDS/NIMH U01NS113252 awarded to NS.

