## SupplementalText for "A multimodal approach for visualization and identification of electrophysiological cell types *in vivo*"

### Supplementary Figures

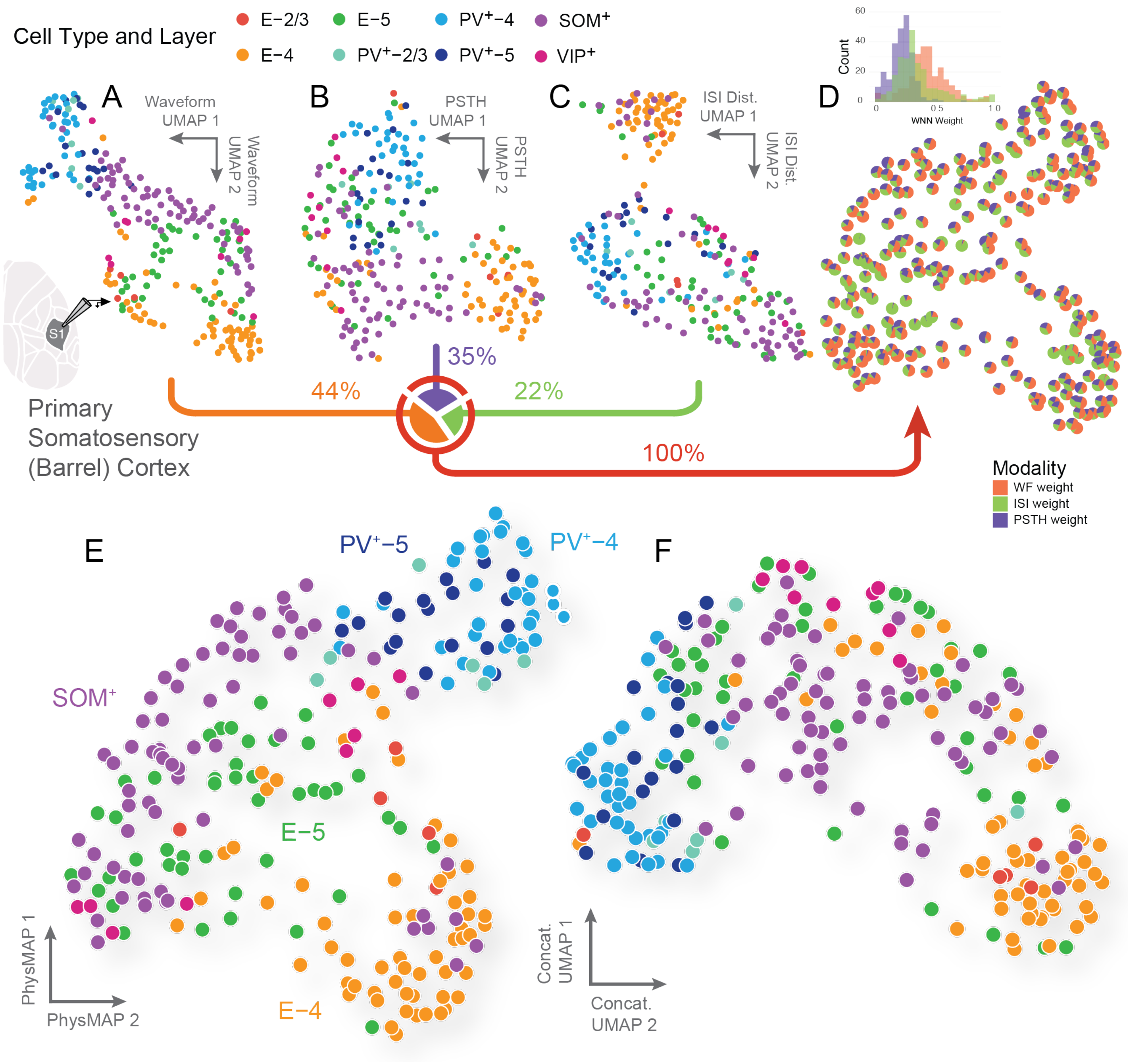


Figure S1: **Multi-modal combination of three physiological properties also leads to structured representations that align with cell types**. (**A**) UMAP on normalized average waveform shapes of neurons (n = 2). Each unit is colored according to one of eight ground truth cell type and layer listed above. (**B**) UMAP on ISI distributions of the same neurons in (A). (**C**) UMAP on the PSTH firing rate distributions from each neuron in response to a deflection of a whisker during whisking. (**D**) Modal contributions to each unit which are the sum of nearest neighbor edge modality weights ($\beta_{\text{Modality}}$ in Fig. A.13J) and are shown as proportions of each pie chart. (**E**) The multimodal representation via the weighted nearest neighbors approach[38].(**F**) All three modalities in the dataset from **(author?)**[32] were concatenated into a single data vector per unit and passed into UMAP. This represents each modality in unweighted combination (as opposed to the WNN approach in PhysMAP).

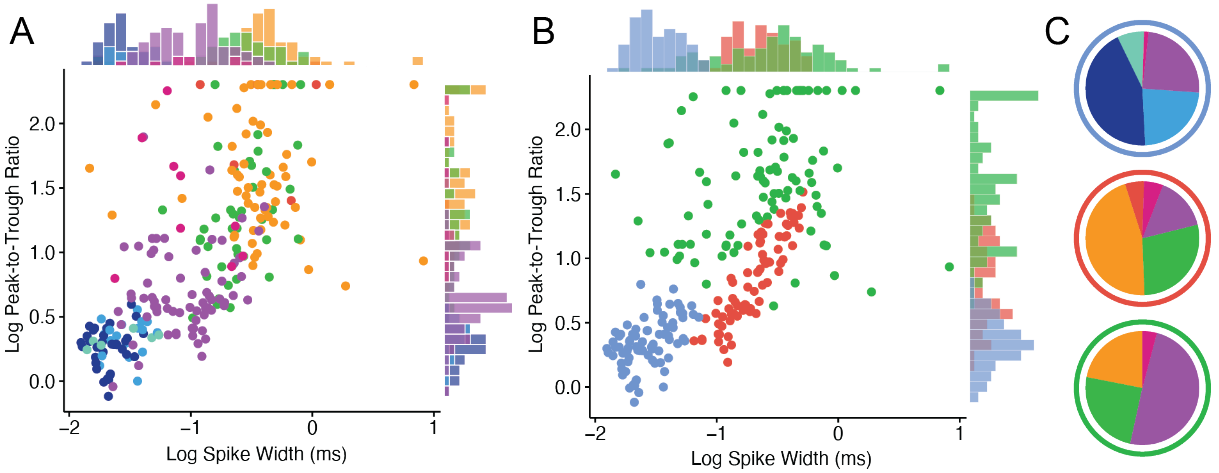


Figure S2: **Waveform shape metrics offer intuition but are less well-connected to cell types**: (**A**) The neurons in Fig. 1 now in a scatter plot according to the log transformation of spike width (in ms) and peak-to-trough ratio. Histograms for each feature metric are shown on the opposite marginals. This is approximately a reproduction of the plot in Fig. 1E of[103]. Note that some peak-to-trough ratios were very large and so are truncated to a maximum value. (**B**) A Gaussian mixture model (GMM) clustering at optimal Bayesian information criteria showing three clusters with densities on the opposite marginals. (**C**) Each GMM cluster is broken down into their constituent cell types and displayed proportionally.

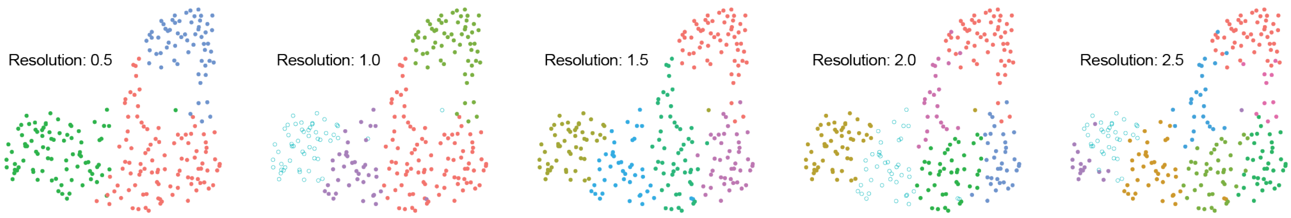


Figure S3: **Sample Leiden clustering visualizations of PhysMAP representations at different resolution parameters**. Each panel shows the same dataset clustered with the Leiden algorithm at different resolution values (0.5, 1.0, 1.5, 2.0, and 2.5). Points are colored according to the cluster assignments determined by the algorithm. As the resolution parameter increases from left to right, more clusters are detected, capturing finer-grained structure in the data. This visualization demonstrates how the selection of the resolution parameter affects the granularity of cell type identification in the PhysMAP embedding space. Higher resolution values result in the detection of more potential cell subtypes, while lower values identify broader cell type categories. These clustering results were used to calculate the MARI scores comparing clustering outcomes with ground truth cell type labels as shown in the main text Fig. 2C.

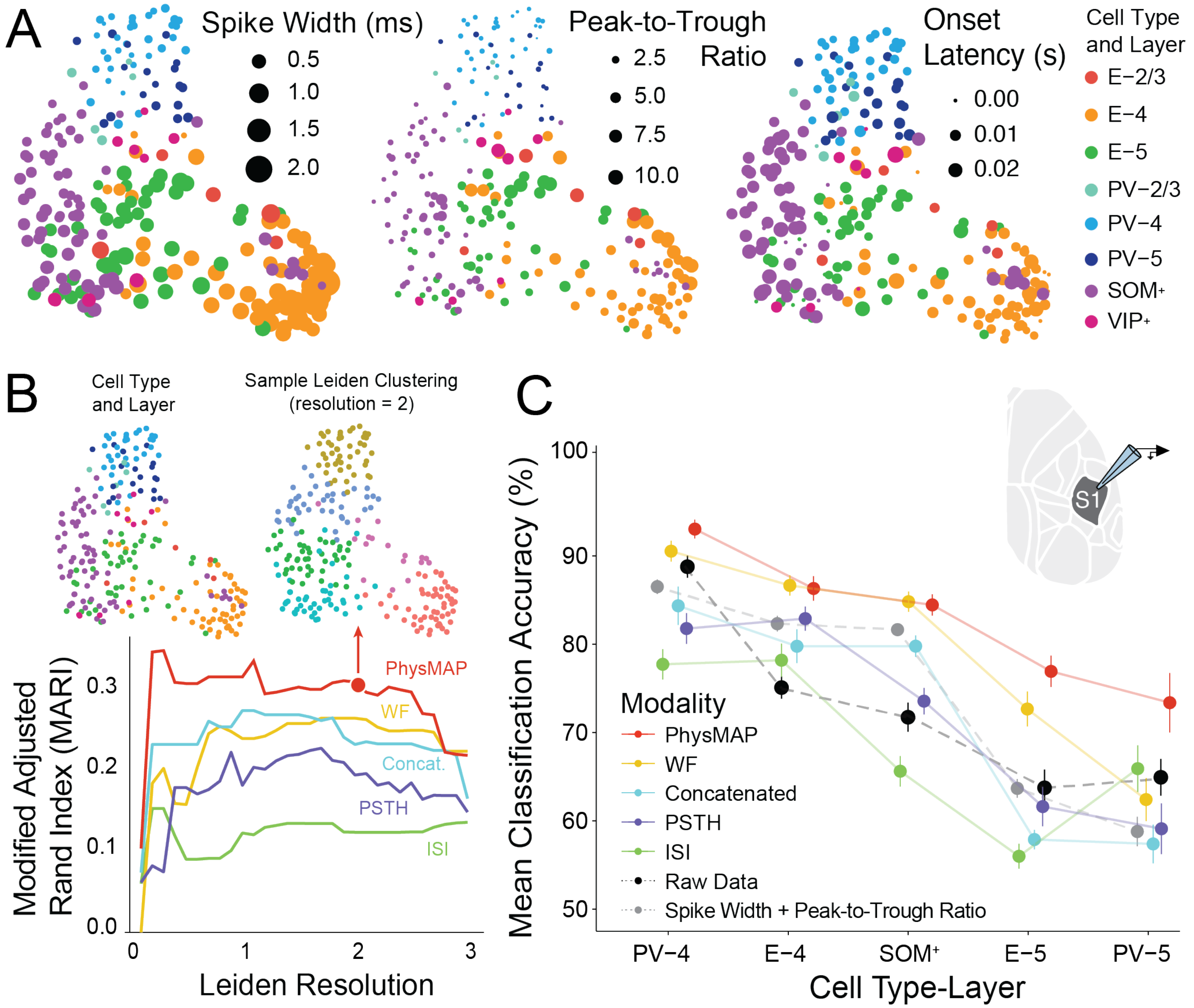


Figure S4: **PhysMAP identifies cell types from juxtacellular recordings better than any modality alone.** (**A**) A gradient boosted tree model (GBM) classifier was trained with 5-fold cross-validation on the 10-dimensional projection of each modality's UMAP graph individually or on the 10-dim. projection of the multimodal WNN graph. The same classifier was also trained directly on the full data (that is, without constructing a UMAP graph and projecting it), on the 10-dim. projection of the UMAP graph of the data concatenated into a single feature, and on the two derived waveform metrics (spike width and peak-to-trough ratio). The balanced accuracy performance of this classifier (mean $\pm$ S.E.M.) on held-out data for each modality, combined modalities, and derived metrics is shown for the five cell type classes with over 10 units each. (**B, top**) The PhysMAP projection of neurons with their ground truth cell type and layer (left) next to an example Leiden clustering[48] with resolution parameter set to 2. (**B, bottom**) Leiden clustering is applied to the UMAP graphs of each modality alone (waveform [WF], ISI dist., and PSTH), in weighted (PhysMAP), or unweighted combination (concat.) and shown with the associated modified adjusted Rand index (MARI;[49]) calculated across a range of resolution parameter values from 0.1 to 3.0 in 0.1 step increments. Arrow marker indicates the clustering on PhysMAP used above in B. Waveform metrics and raw data were omitted in this analysis because these do not yield a high-dimensional UMAP graph. (**C**) The spike width, peak-to-trough ratio, and spiking onset latency for each neuron are shown via log transformed marker size under PhysMAP's two-dimensional projection.

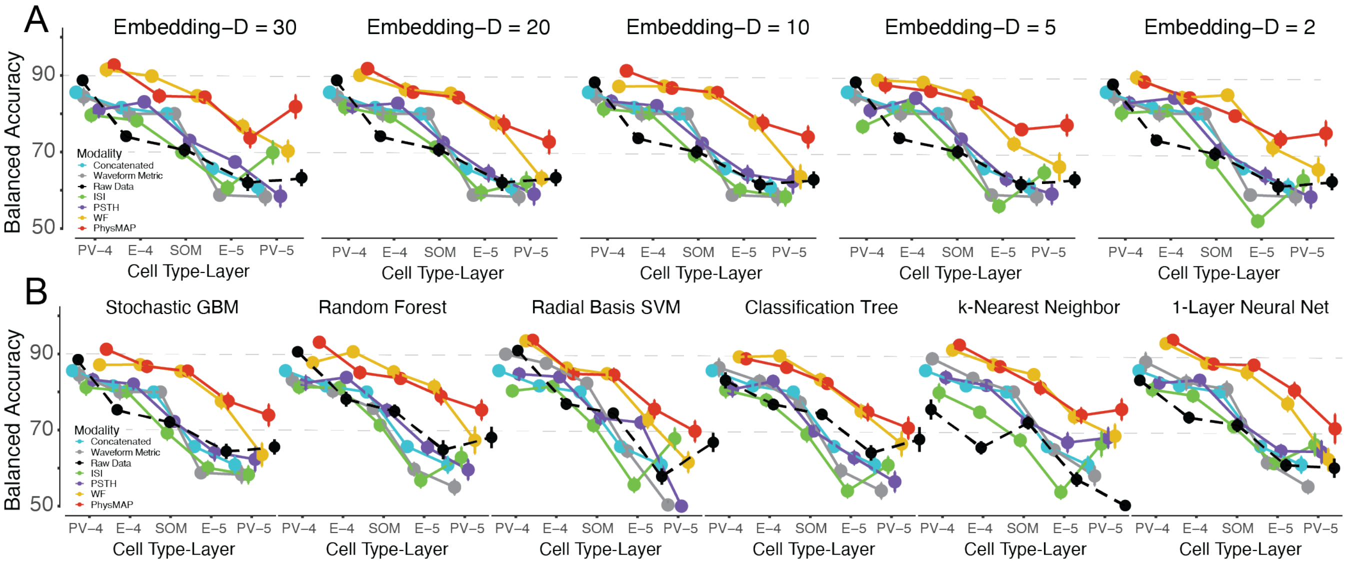


Figure S5: **Classifier performance is not affected by graph embedding dimensionality or choice of classifier**: (**A**) The embedding dimensionality of the WNN graph was varied between 30 and 2 dimensions and used for training the same GBM classifier as in Fig. 2F which was trained on embedding dimension of 10 with five-fold cross-validation. The mean balanced accuracy ($\pm$ SEM) across the five cell types with more than 10 examples is shown. (**B**) With a WNN graph embedding dimension of 20, six different classifiers (with default *caret* settings) were trained to identify the five cell types in Fig. 2F (which was the same here as the stochastic GBM). These plots are once again shown with mean balanced accuracy ($\pm$ SEM) after five-fold cross-validation.

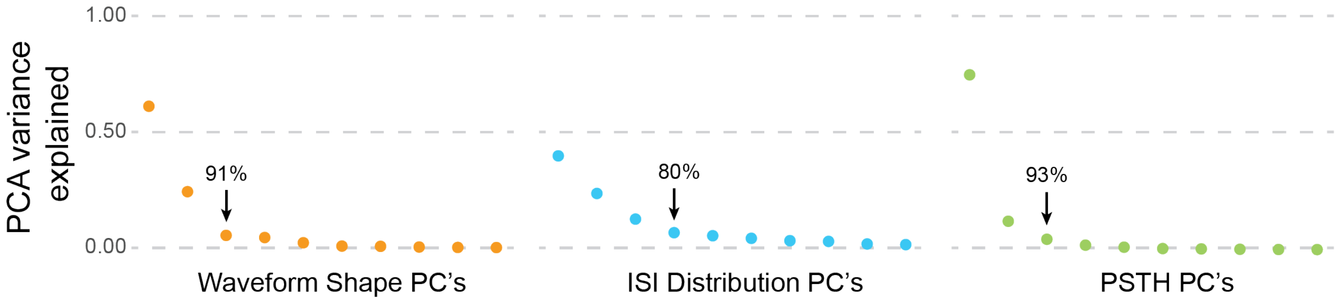


Figure S6: **Principal component analysis gives an estimate of linear intrinsic dimensionality**: Principal component analysis (PCA) was applied to the data from each modality in Fig. 1A including PSTH (not used) and the per-component variance explained calculated for each. For each modality, the scree plot shows the variance explained for each principal component. An arrow on each plot demarcates the "elbow" with cumulative percent variance explained listed above.

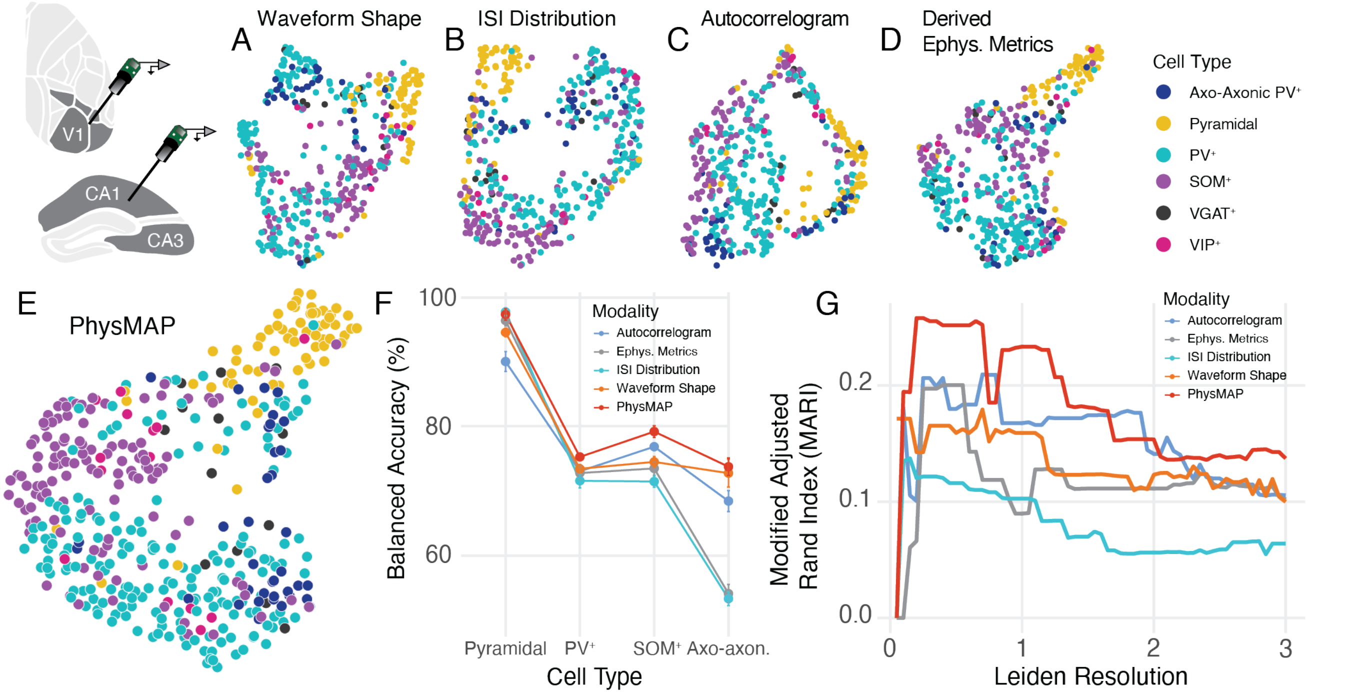


Figure S7: **PhysMAP is able to identify cell types across areas and labs**. (**A** - **D**) Across a pooled dataset of visual cortex and hippocampal extracellular recordings from CellExplorer, we show 2-D UMAP visualizations applied to waveform shape, inter-spike interval distribution, autocorrelogram, and thirteen various derived electrophysiological metrics respectively. Cell type identifications were obtained by opto-genetic tagging. (**E**) The previous four modalities are combined into a unified PhysMAP representation and visualized in 2D. (**F**) A gradient boosted tree model (GBM) classifier with 5-fold cross-validation was trained on the 10-dimensional projection of each modality's high-dimensional UMAP graph or on PhysMAP's multimodal WNN graph. The balanced accuracy for the classifier is shown for each modality and cell type that contained more than 25 examples. (**G**) Leiden clustering is applied to each modality alone and to PhysMAP and the corresponding modified adjusted Rand indices are calculated across a range of resolution parameter values from 0.1 to 3.0 in 0.1 step increments. All cell type labels were included in this calculation.

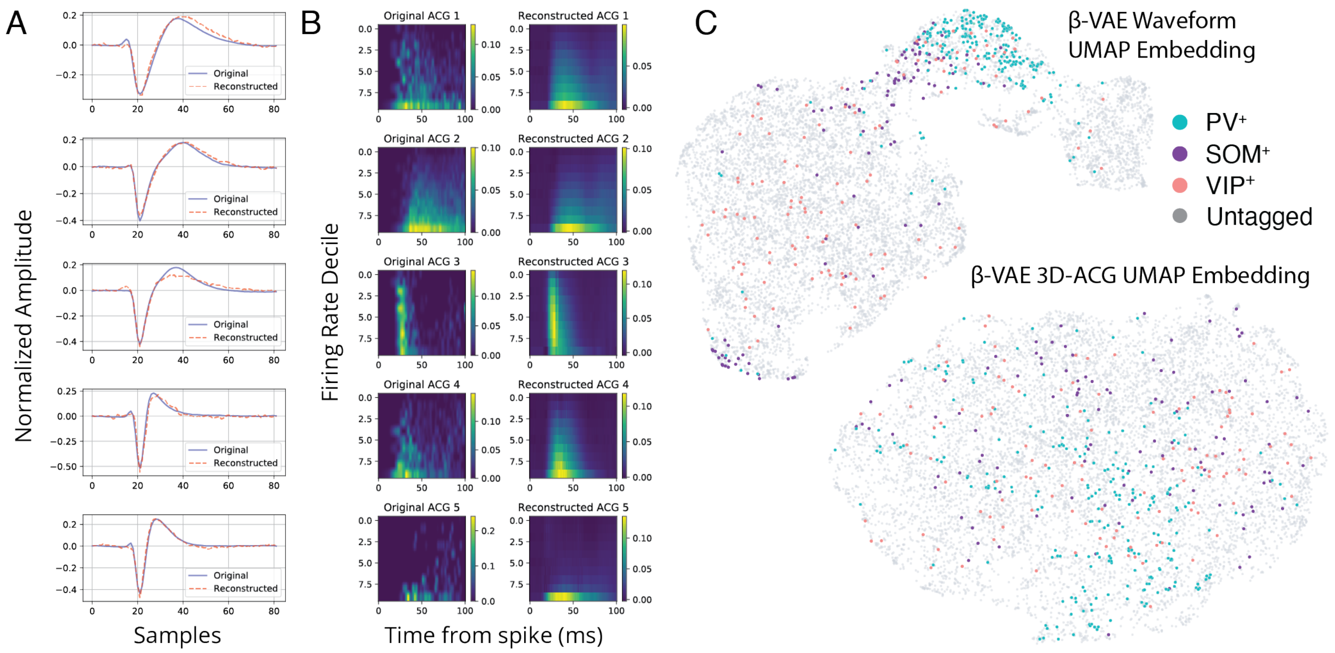


Figure S8: $\beta$-**variational autoencoder reconstructions for each modality along with embedding visualizations** (**A**) Five sample normalized average waveform shapes (original; blue line) along with their VAE reconstructions (dashed red line) from the Ultras dataset. (**B**) 3D-autocorrelograms for the same five units with original input data (left column) and output reconstructions (right column). (**C**) Visualizations of each VAE's latent embeddings for waveform shape (**top**) and 3D-ACG (**bottom**). Each data point is color coded according to its optotagged identity.

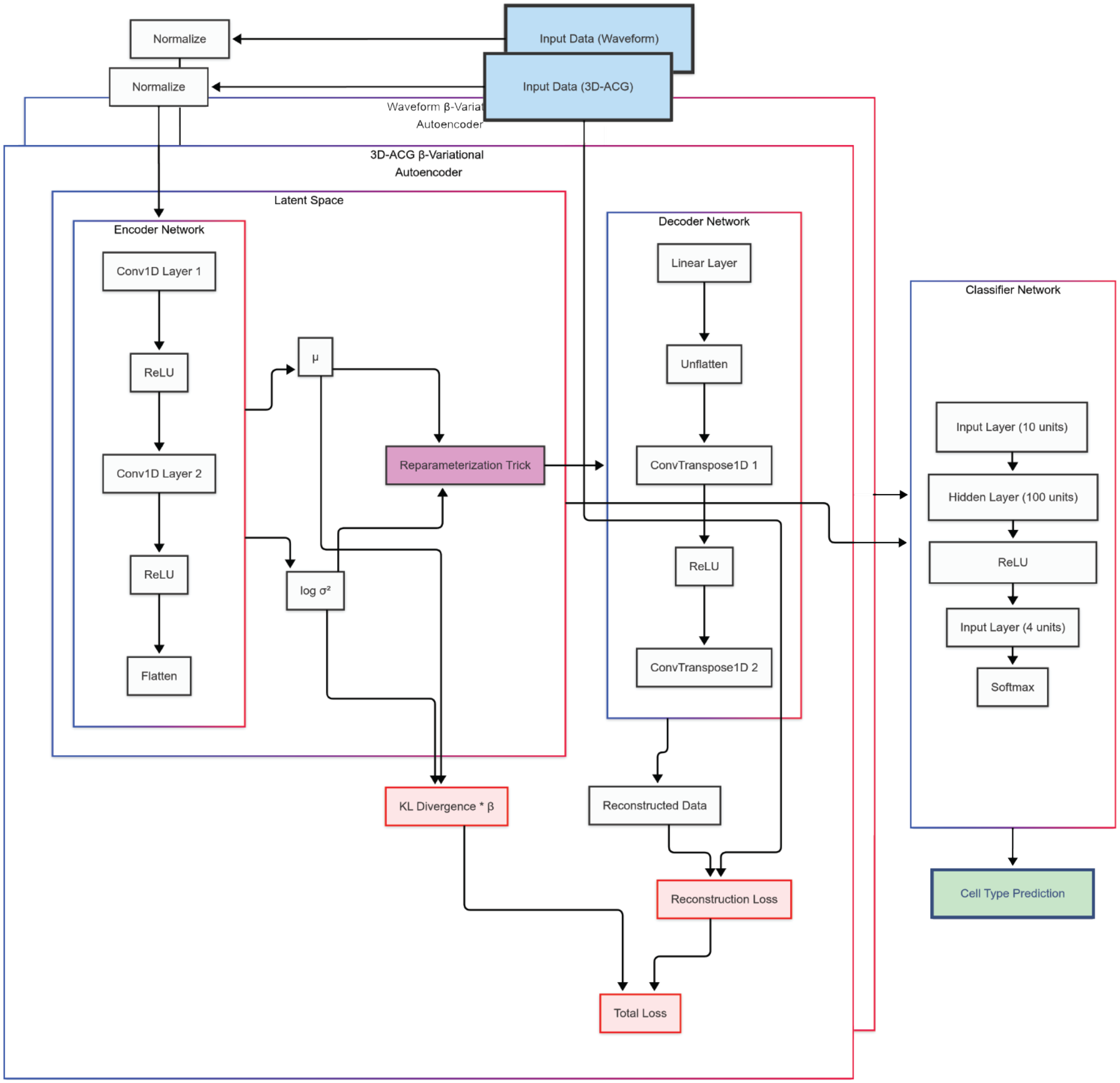


Figure S9: **The dual $\beta$-variational autoencoder architecture**. Two identically structured VAE's are used with each encoding one of the two modalities used (waveform shape and 3D-ACG) with individually-tuned $\beta$ parameter. Each modality is normalized and passed into separate encoder networks consisting of two convolutional layers with ReLU nonlinearities after each. The reparametrization trick is applied the the compressed representation which is then decoded by a decoder network consisting similarly of two convolutional layers and associated ReLU nonlinearities. This yields a reconstruction which is compared against the original data via the ELBO (evidence lower bound) penalty minimizing Kullback-Liebler divergence and reconstruction loss. After training the $\beta$-VAEs, the both pass their latent embeddings to a single multi-layer perceptron (consisting of a single hidden layer with ReLU nonlinearity).

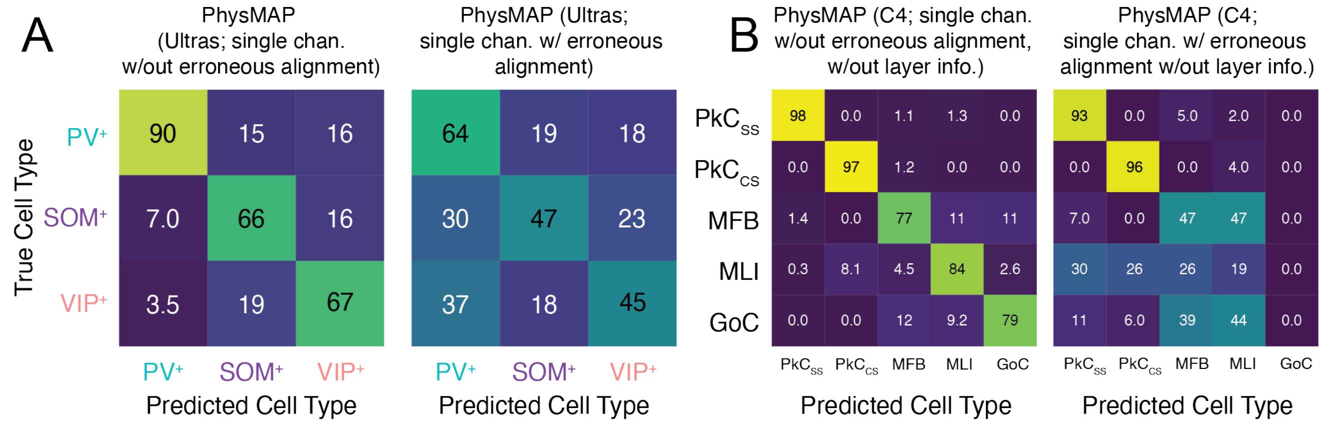


Figure S10: **PhysMAP without erroneous landmark alignment provides superior performance across multiple datasets.** (**A**) Confusion matrices showing classification performance with a gradient-boosted tree model for three inhibitory neuron types ($PV^+$, $SOM^+$, and $VIP^+$) in the Ultras dataset[57] using single-channel PhysMAP without alignment (**left**) and with erroneous alignment (**right**). Values represent percentages of true cell types classified correctly as each predicted type. (**B**) Confusion matrices showing classification performance for five cerebellar cell types (Purkinje cell simple spikes [$PkC_{SS}$], Purkinje cell complex spikes [$PkC_{CS}$], mossy fiber bundles [MFB], molecular layer interneurons [MLI], and Golgi cells [GoC]) in the C4 dataset[63] using single-channel PhysMAP without alignment and without layer information (**left**) and with alignment but without layer information (**right**). Values represent percentages of true cell types correctly classified as each predicted type.

### Appendix A. Multi-modal analysis of extracellular electrophysiology with PhysMAP

Our PhysMAP approach uses a weighted-nearest neighbor graph combination solution developed in multiomics for intelligently integrating multiple modalities (transcriptomic, epigenomic, and proteomic data). This is in order to find latent cell type structures that transcend any single modality[38]. This multimodal approach is common in machine learning and provably outperforms unimodal approaches[104]. As in our previous WaveMAP approach[25,42], PhysMAP uses the average normalized data for each single unit. Unlike WaveMAP, PhysMAP looks at arbitrarily many modalities beyond simply waveform shape. PhysMAP then finds shared underlying structure by combining these modalities via a weighted nearest neighbor (WNN) graph. After assessing all modalities for all cells, this method unifies all modality-specific graphs by weighing each according to their informativeness on a per-unit basis. This graph is then projected into two dimensions for visualization of multimodal structure, but its higher-dimensional graph can be used for cell type identification. In Fig. A.13, we show how this WNN graph is constructed in the simple two-modality case combining waveform shape and ISI distribution. What follows here is a description of these steps followed by the motivation for each. This methodology is described in greater detail in Methods: Weighted Nearest Neighbors.

1. **Within-Modal Affinity**: Within the waveform space (Fig. A.13A), we select a single neuron (blue sphere in Fig. A.13B), identify its $k$-nearest neighbors (in this example, $k$=5; red spheres in Fig. A.13B), and average them to predict the waveform of said neuron (dashed red circle in Fig. A.13C). We then calculate the *within-modal affinity* by passing the neuron's actual waveform and its predicted waveform into a modified UMAP distance kernel (Fig. A.13D, numerator), and thus a measure of how well a neuron's waveform is predicted by its neighbors in waveform-space. In the third step, we will show how this will be useful when determining which modalities to "trust" more in the form of an "affinity ratio".
2. **Cross-Modal Affinity**: Next, we calculate a *cross-modal affinity* for the ISI distribution modality (Fig. A.13E). Using the same neuron and its same neighbors in Fig. A.13B, but now in ISI-space where each dimension is a time point along an ISI distribution curve (Fig. A.13F), we calculate a predicted ISI distribution for this neuron by averaging (dashed red circle in Fig. A.13G). We calculate affinity again by

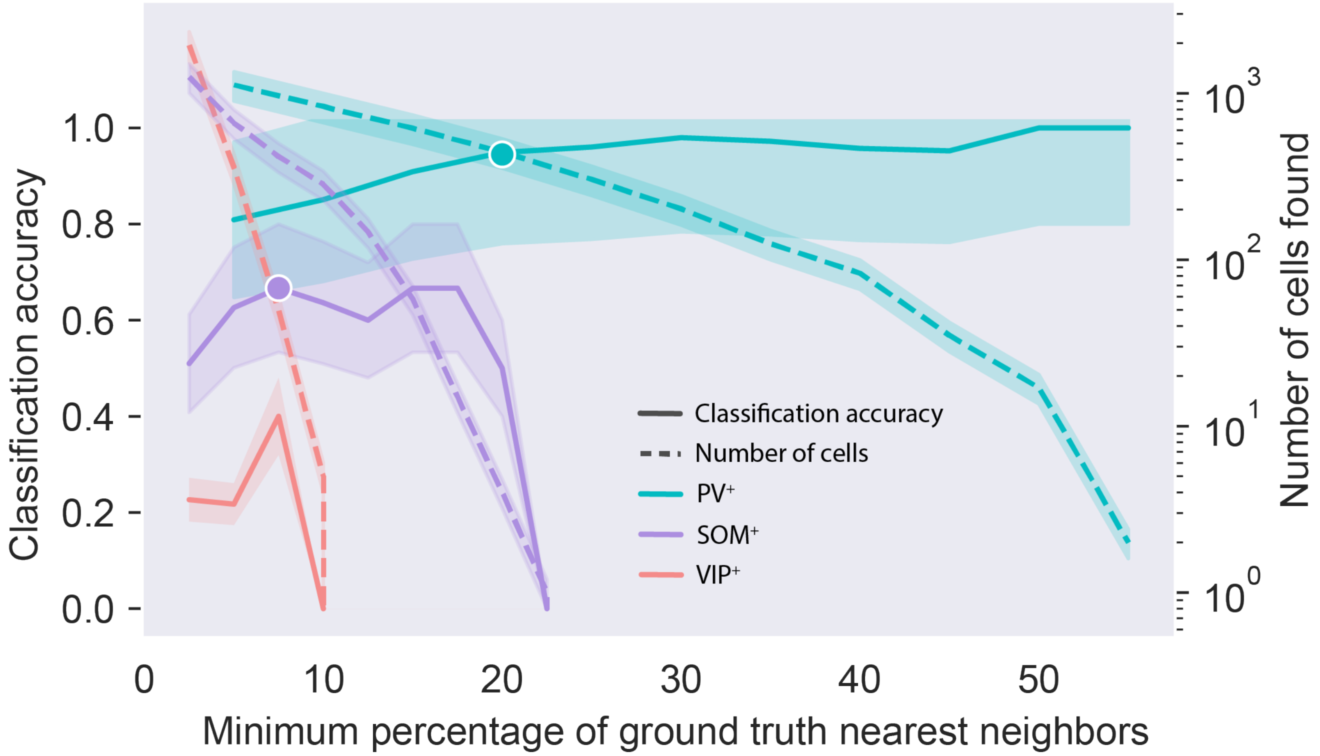


Figure S11: **Increasing the percentage of neighbors sharing a label to make a prediction is positively correlated with classification accuracy but negatively correlated with yield**. The plot shows the relationship between classification accuracy (solid lines, left y-axis) and number of cells identified (dashed lines, right logarithmic y-axis) as a function of the minimum percentage of nearest neighbors on the high-dimensional PhysMAP graph required for cell type identification. Data is shown for three inhibitory neuron types: $PV^+$ (teal), $SOM^+$ (purple), and $VIP^+$ (red). White circles indicate the selected threshold values used for cell type identification in the main analysis for $PV^+$ and $SOM^+$ neurons.

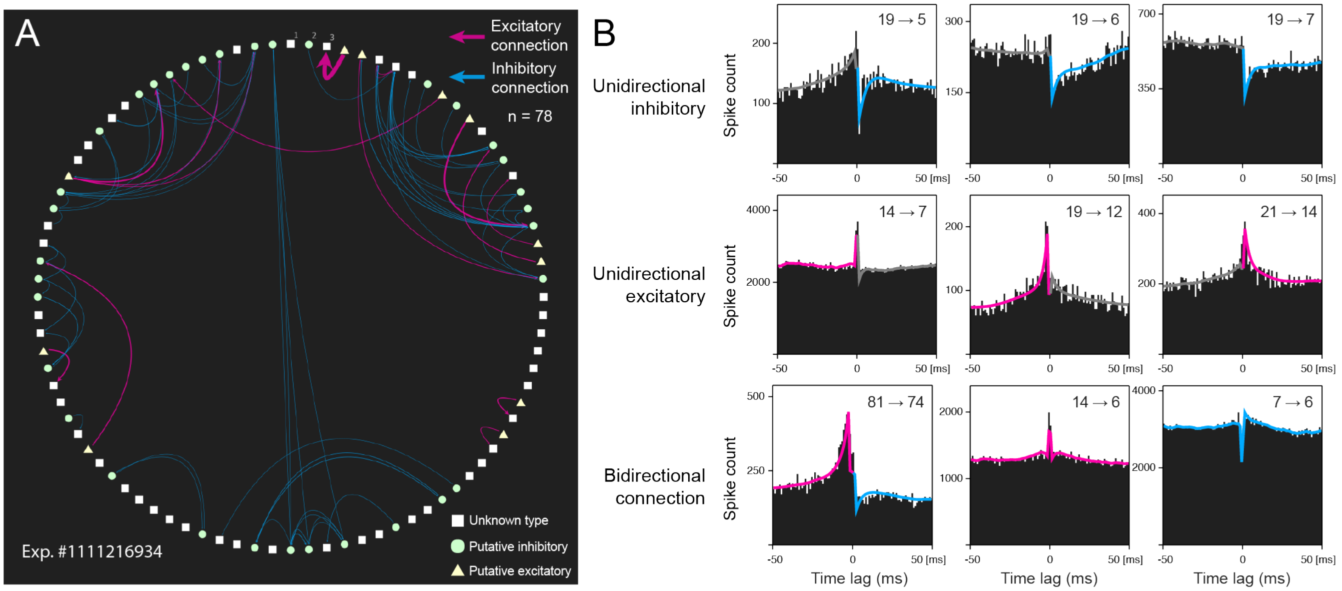


Figure S12: **Putative monosynaptic connectivity network and cross-correlograms of neurons in a single Visual Behavior experiment** (**A**) Connectivity diagram showing putative connections between neurons recorded in experiment #1111216934 of the Visual Behavior dataset. Nodes represent individual neurons arranged in a circle, with putative inhibitory neurons (green circles), putative excitatory neurons (yellow triangles), and neurons of unknown type (white squares). Directional putative monosynaptic connections between neurons are shown as curved arrows: excitatory connections in magenta and inhibitory connections in blue. The network contains 78 total connections. These were the interactions deemed significant ($p < 0.0001$) after applying the GLMCC approach with default parameters[76] to spike times from the entire session. (**B**) Example cross-correlograms showing spike-timing relationships between connected neuron pairs. Each panel displays spike count (y-axis) relative to reference spike times expressed as lags (x-axis, in milliseconds) with neuron IDs indicated as source→target. Cross-correlograms are grouped by connection type: unidirectional inhibitory (**top row**, blue trace overlay), unidirectional excitatory (**middle row**, magenta trace overlay), and bidirectional connections (**bottom row**, with both inhibitory and excitatory components).

passing the neuron's true ISI distribution and it's predicted ISI distribution into a modified UMAP distance kernel (denominator of Fig. A.13D), providing a measure of how well one modality in the space of another's space is predictive of a neuron's properties (using the same neighbors as to facilitate comparison).

3. **Affinity Ratio**: We define the per-unit *waveform affinity ratio*, $\mathsf{S}_{\mathsf{Wave}}(i)$, (fraction in Fig. A.13D) as the ratio of this neuron's waveform *within-modal affinity* divided by its ISI dist. *cross-modal affinity*. Conversely, the ISI distribution per-unit affinity ratio $\mathsf{S}_{\mathsf{ISI}}(i)$ is also obtained (not shown) by considering the ISI distribution-space as the within-modality and the waveform shape-space as the cross-modality. This ratio compares how well one modality versus another is predictive of a neuron's properties and will be used to determine how much to weigh each modality for this neuron. In the case that there are more than two modalities, the denominator sums over all cross-modality distances.
4. **Modality Weight:** We then calculate the weight associated with a unit for a particular modality ($\beta_{\mathsf{Modality}}(i)$) as the ratio of the exponentiated affinity ratio for a modality divided by the sum of exponentiated affinity ratios over all modalities (Fig. A.13H; left equation for waveforms, right equation for ISI dists.). Thus for each neuron, each modality is differentially weighted according to how well the cell's properties are predicted in each.
5. **Pair-Wise Unit Distances**: The distances between every pair of points (in every modality) is also passed into the same modified UMAP distance metric for construction of each modality-specific graph before re-weighting (Fig. A.13I). For example, the edge between a pair of units $i$ and $j$ in waveform shape-space is $\theta_{\mathsf{Wave}}(i, j)$. In this way, subsequent calculations use UMAP's notion of distance which has been shown to better capture underlying low-dimensional manifold structure[37].
6. **WNN Construction and Visualization**: Finally, we calculate a connectivity matrix of new pair-wise edges (distances) by taking the weighted linear combination of edges in the waveform space and ISI space with the pair-wise distances and affinity weights calculated in the previous steps (Fig. A.13J). The final WNN graph is derived from this matrix by using *k*-nearest neighbors algorithm with *k* set to a large number (here $k = 200$). Using UMAP's force-directed graph layout procedure, we then visualize the high-dimensional multimodal structure by projecting the graph into two dimensions. The unprojected graph is used for identification.

These steps proceed similarly in the case of three or more modalities with affinity ratios calculated for each modality against every other. The summations in the denominators of Fig. A.13H and the weighted average of Fig. A.13J expand to include these other modalities as well. In the next sections, we use this PhysMAP approach and show how it can combine multiple modalities to delineate cell types in three different datasets.

### Appendix B. Bias due to unsupervised transformation before data splitting is negligible

In its current form, the WNN construction doesn't admit a function that can be reused to classify untagged cells once trained—all cells to be classified must be included in the transformation itself. While this is not an issue as PhysMAP is highly performant if needing to be rerun for application to new data, it does mean our cross-validation procedure applied a test-train split *after* a data-dependent transformation was applied. This can result in a more unconventional form of data leakage wherein test information leaks into the training dataset via its effect on the computed unsupervised transformation itself[64]. However, this effect is usually pessimistic i.e., training accuracy underperforms test accuracy. Furthermore, this is usually only an issue for datasets numbering less than around 100 data points[64]. To check that we are not overestimating our performance on held-out data, we adopt the procedure of[64] which entails transforming the entire dataset with PhysMAP but then conducting the test-train splitting on only a subset of the data; the performance on the test set (which is still within the data subset) is regarded as the "validation error." The classifier trained on the subset is then evaluated against the held-out data and this is referred to as the "generalization error." In this way, by examining the difference between generalization and validation errors, it is evaluated whether or not the particular characteristics of the dataset and transformation yield an optimistic or pessimistic estimate of classifier generalization due to data leakage and to what degree. The data subset proportion is increased and this process is reevaluated to evaluate

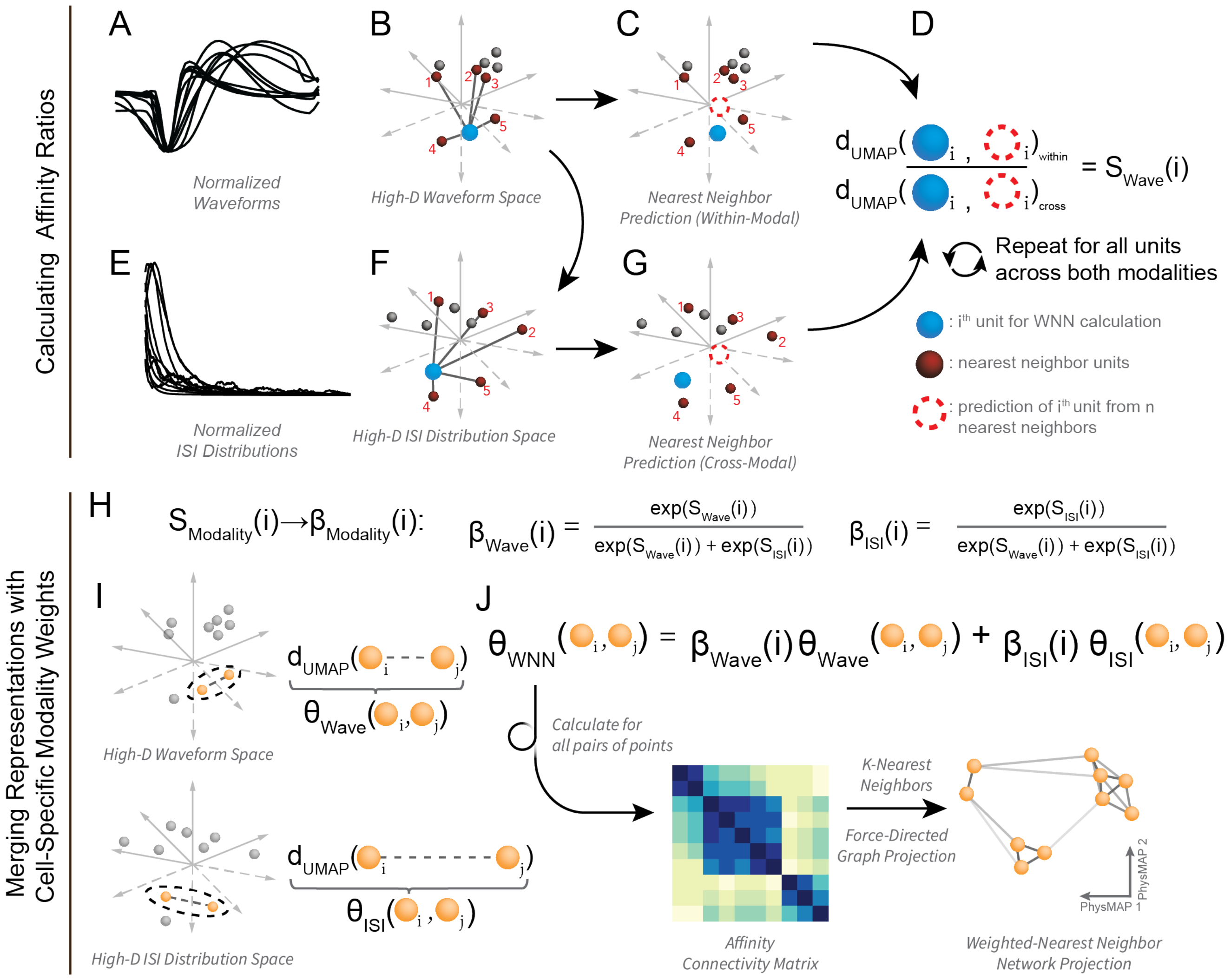


Figure A.13: **Schematic of PhysMAP and weighted-nearest neighbors algorithm**. (**A**) A sample of normalized average single unit extracellular action potential waveforms. (**B**) Waveforms in (A) are shown in high-dimensional space with each axis pertaining to each time point along the waveform's trace. For a sample unit (blue sphere), its nearest neighbor waveforms are highlighted (numbered red spheres). (**C**) The nearest neighbor waveforms are averaged to generate a prediction of the original waveform (dashed red circle). (**D**) The numerator of the waveform affinity ratio ($S_{\text{Wave}}(i)$), the "within-modal affinity", is the difference between the original neuron's average waveform and its nearest neighbor prediction. This "reconstruction error" distance is passed through a modified UMAP distance kernel. (**E**) Normalized ISI distributions drawn from the same neurons that produced the waveforms in (A). (**F**) The same neuron in (B; blue sphere) and its same nearest neighbors in waveform space now in ISI-space. (**G**) A prediction is made for the sample unit in ISI-space (red dashed circle). The difference between these is the "cross-modal affinity". The ratio of within- and cross-modal affinities form the per-unit waveform affinity ratio ($S_{\text{Wave}}(i)$). (**H**) Per-unit affinity ratios ($S_{\text{Modality}}(i)$) for each modality are converted into per-unit modality weights ($\beta_{\text{Wave}}(i)$ and $\beta_{\text{ISI}}(i)$). (**I, top**) The difference between a sample pair of points in high-dimensional waveform space is another type of "affinity" ($\theta_{\text{Wave}}(i,j)$) passed into UMAP's modified distance kernel. (**I, bottom**) The same pair of points as in (I, top) with the modified UMAP distance between them in ISI-space ($\theta_{\text{ISI}}(i,j)$). (**J**) Weighted affinities ($\theta_{\text{WNN}}$) as a weighted combination of unimodal affinities shown also as an pairwise affinity matrix. Two-dimensional UMAP projection of the weighted *k*-nearest neighbors of the affinity connectivity matrix. Note that affinity distances $\theta_{\text{WNN}}$ are not necessarily symmetric from as shown in the heatplot i.e., it is not always true that $\theta_{\text{WNN}}(i,j) = \theta_{\text{WNN}}(j,i)$.

this potential bias as data size increases. From[64] it is shown that, for all but the most pathological cases, this difference trends to zero quickly and classifier performance plateaus as sample size increases; we show here that this too is the case for our dataset.

In Fig. S13A, to assess the magnitude and direction of any potential bias, we conducted this procedure while increasing the proportion of the full dataset. Visually, validation accuracy ($1 -$ validation error) increased in concert with generalization accuracy ($1 -$ generalization error) but seemed to be nearly equivalent at all data proportions beginning at around 80 data points. To evaluate the direction of any potential bias, we plotted a rolling average of the difference between validation and generalization at all data proportions (Fig. S13B). We found that the direction of the bias was consistently negative across all data proportions i.e., the bias was pessimistic. To estimate the magnitude of this bias, we performed a kernel density estimation across each data proportion and examined the median relative to zero (Fig. S13C). While the median of this density estimation was less than zero, it was not significant (Wilcoxon signed-rank test). Thus, we are confident that even though we are conducting an unsupervised data transformation before data splitting, if a bias exists, it is negligible.

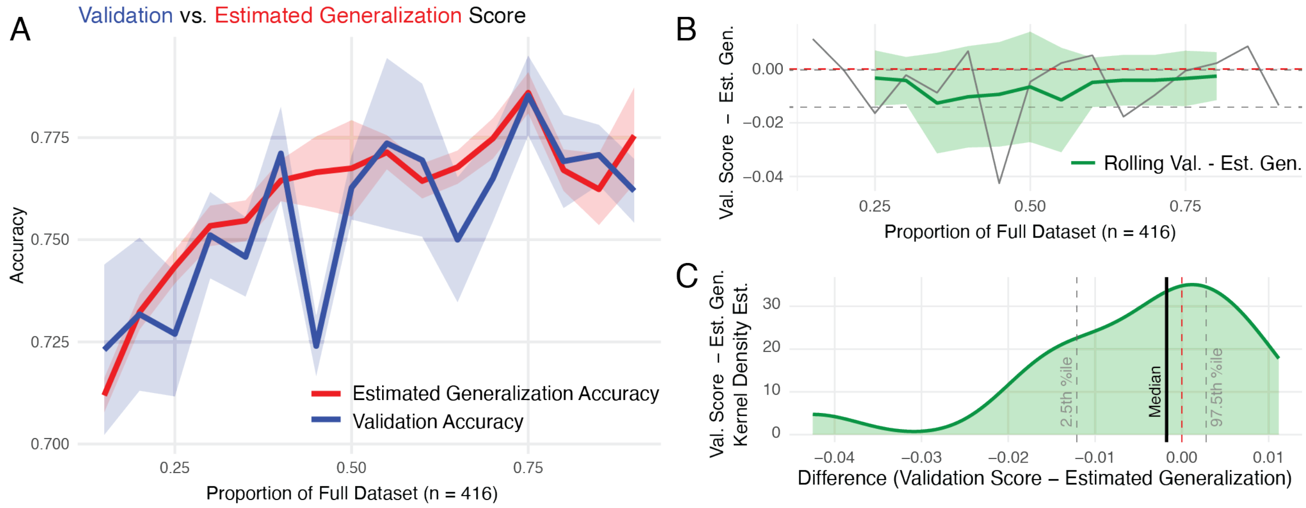


Figure B.14: **Simulation shows negligible potential bias from unsupervised transformation before data splitting**. (**A**) Two accuracy metrics plotted against the proportion of the full dataset (n=416): validation accuracy (blue line) and estimated generalization accuracy (red line), both with shaded SEM. These were calculated at each proportion over 25 runs with randomized initial. (**B**) shows the difference between validation score and estimated generalization score (green line with shaded SEM) across dataset proportions from 0.25 to 0.75. (**C**) displays a Gaussian kernel density estimation of the difference values (validation score minus estimated generalization), with vertical gray dashed lines marking 2.5th percentile and 97.5th percentile.
